# CpG-induced regions associated compositional stratification of human essential proteins through mathematical genomics

**DOI:** 10.1101/2025.11.21.689746

**Authors:** Shipra Singh, Kharerin Hungyo, Sk. Sarif Hassan, Vladimir N. Uversky

## Abstract

CpG islands are genomic regions enriched in cytosine–phosphate–guanine dinucleotides, typically associated with gene promoters and regulatory elements. While their role in transcriptional regulation is well established, their influence on protein sequence composition remains underexplored. In this study, 3,222 essential proteins from *Homo sapiens* were analyzed to investigate the impact of CpG island architecture on amino acid usage, sequence complexity, and chromosomal distribution. Codons containing both cytosine and guanine were mapped to five CpG-induced amino acids, and their relative abundance was quantified across proteins. CpG-induced regions were identified using a sliding window approach, and metrics such as average CpG density, longest CpG-induced region length, and sequence coverage were computed. Motif composition within these regions was characterized using polarity and charge-based indices. Clustering revealed consistent bipartite subgroup structures, indicating compositional stratification among essential proteins. Shannon entropy quantified compositional complexity, revealing a significant shift between clusters (*p* = 5. × 50 10^−145^). Cluster 1 proteins showed lower entropy and greater heterogeneity, whereas Cluster 2 showed higher entropy and tighter distributions, indicating distinct regimes of variability. Additionally, we examined CpG island overlap within coding exons of essential genes and found that CpG-depleted proteins are predominantly encoded by genes with minimal CGI coverage, whereas CpG-induced proteins span both low- and high-overlap classes. CGI-associated CpGs were consistently enriched toward the 5′ end of essential genes, indicating a positional bias in CpG distribution. These findings establish a quantitative framework linking CpG island distribution to proteomic architecture and offer a scalable strategy for motif annotation, epigenetic modeling, and functional stratification. Future applications include predictive modeling of protein function, disease association, and regulatory dynamics.

## 1. Introduction

CpG islands (CGIs) are short stretches of DNA characterized by high GC content and an elevated frequency of CpG dinucleotides relative to the rest of the genome [1, 2, 3]. Approximately 45,000 CpG islands are estimated in the human genome, with the majority located near gene promoters and transcription start sites, often marking regions of active or potential transcription [4, 5]. Unlike the bulk genome, which is depleted of CpG dinucleotides due to the mutational conversion of methylated cytosines to thymines, CpG islands remain largely unmethylated and are evolutionarily conserved as key regulatory elements [6, 7, 8]. Their unmethylated status facilitates transcription factor binding and the establishment of open chromatin configurations, thereby maintaining gene accessibility [9]. Aberrant methylation of CpG islands, on the other hand, leads to transcriptional silencing and has been implicated in various biological processes such as X-chromosome inactivation, genomic imprinting, aging, and tumorigenesis [10, 11, 12].

The chromosomal distribution of CpG islands is non-random; they are predominantly enriched in R-band, GC-rich, and early-replicating regions of the genome that coincide with high gene density [13, 14]. Functionally, CpG islands act as epigenetic hotspots that integrate DNA sequence composition with chromatin states [6, 7, 15]. Promoter-associated CpG islands are typically long, GC-rich, and stably hypomethylated, supporting constitutive expression of housekeeping genes, while those located in intragenic or intergenic regions often exhibit tissue-specific or developmentally dynamic methylation patterns [7, 16]. Recent epigenomic analyses have also revealed that CpG islands recruit histone-modifying enzymes through the recognition of unmethylated CpG motifs, thereby establishing unique chromatin architectures that distinguish these regions from the surrounding genome [17, 18, 19]. Thus, CpG islands serve not only as genomic markers of gene regulatory regions but also as functional platforms that couple DNA methylation status, transcriptional control, and chromatin organization [18, 20, 21].

Advances in computational and genome-wide mapping techniques have significantly enhanced the identification and characterization of CpG islands, enabling integrative analyses linking CpG density, methylation profiles, and transcriptional activity [6, 22]. Emerging evidence suggests that the nucleotide composition surrounding CpG islands, particularly in promoter-associated genes, may influence codon bias and amino acid composition within coding regions, thereby connecting epigenetic regulation to protein-level characteristics [23, 24]. Understanding the functional implications of CpG island distribution and their downstream effects on gene and protein properties offers a promising avenue to explore the evolutionary and regulatory principles that underlie the expression and stability of essential human proteins.

Essential genes and their encoded proteins represent the fundamental molecular framework required for cell survival, growth, and genomic stability [25, 26, 27, 28]. These genes are typically characterized by evolutionary conservation, ubiquitous expression, and tight regulatory control, often mediated through CpG-rich promoters [7, 29]. Given the centrality of CpG islands in transcriptional regulation, methylation dynamics, and chromatin accessibility, their distribution and composition within essential genes may have profound implications for maintaining expression fidelity and functional robustness [30, 31]. However, despite extensive cataloging of CpG islands across the human genome, a systematic and quantitative understanding of their occurrence, density, and compositional influence in essential genes remains limited [32]. Exploring the interplay between CpG island architecture and amino acid composition in essential proteins could therefore reveal fundamental principles linking epigenetic regulation, nucleotide composition bias, and protein functionality—providing new insights into how genomic and epigenomic features co-evolve to sustain vital cellular processes in humans.

The chromosomal landscape of essential genes in humans exhibits marked variation in gene density, GC content, and CpG island abundance, suggesting that CpG-driven compositional biases may be unevenly distributed across the genome [32]. By examining CpG-induced amino acid regions (CpG-AARs) across all chromosomes, one can gain insights into how local nucleotide composition constraints translate into amino acid preferences within essential proteins [33]. Because codons containing cytosine and guanine are more frequent in CpG-rich regions, certain amino acids—particularly arginine, alanine, serine, proline, and threonine—tend to cluster within protein segments encoded by CpG-dense DNA. These residues often play crucial roles in structural stability, hydrogen bonding, phosphorylation, and protein–protein interactions, implying that CpG enrichment at the nucleotide level could exert subtle yet systematic influence on protein structure and function.

Comparative analysis of CpG-AARs across human chromosomes provides a quantitative framework to explore how genomic composition and epigenetic organization coalesce to shape proteomic architecture. Chromosomes with higher GC and CpG content, such as chromosomes 17 and 19, are expected to harbor a greater density of CpG-AARs, potentially reflecting their enrichment in essential and housekeeping genes [34, 35, 36]. Conversely, CpG-poor chromosomes may encode essential proteins with fewer or more spatially constrained CpG-AARs, suggesting differential selection pressures across genomic compartments. Mapping and clustering CpG-influenced amino acid regions thus offer a new dimension to understanding how evolutionary, epigenetic, and structural forces jointly maintain the robustness of the human essential proteome.

To systematically investigate the influence of CpG dinucleotide patterns on protein composition, we conducted a suite of quantitative analyses focused on CpG-induced amino acid regions across human essential proteins. By identifying CpG-enriched segments within protein sequences—defined by elevated frequencies of amino acids encoded by CG-containing codons—we computed key metrics such as average CpG density, the length of the longest CpG-induced region, and the fraction of the protein sequence covered by these regions. These metrics were aggregated across chromosomes and clusters to reveal distributional trends. Additionally, motif composition within the longest CpG-enriched region of each protein was quantified using polarity- and charge-based indices, including the ratio of polar to non-polar residues (PolarRatio), and the fractional content of acidic and basic residues. Threshold-based filtering further enabled the identification of proteins with extreme motif enrichment or depletion. Finally, unsupervised clustering based on standardized CpG metrics uncovered consistent bipartite subgroup structures within each chromosome, offering insights into the structural and compositional diversity of CpG-influenced domains in the human proteome.

These analyses revealed distinct patterns of CpG-induced amino acid enrichment and motif composition across human chromosomes, highlighting structural and compositional biases in essential proteins. The identification of cluster-specific and chromosome-specific trends suggests potential regulatory roles of CpG motifs in protein evolution and function.

Emerging evidence suggests that GC-rich and CpG-enriched exonic regions demonstrate distinct chromatin organization, particularly in how DNA interacts with and wraps around nucleosomes. Multiple studies have shown that exons generally exhibit higher nucleosome occupancy than introns, and that this enrichment strongly correlates with GC content and CpG density[37]. Sequence-intrinsic properties of GC- and CpG-rich DNA enhance its affinity for histone octamers, leading to preferential nucleosome positioning over CpG-dense exons[38]. In addition, CpG methylation further modulates nucleosome stability and wrapping, influencing chromatin structure and transcriptional accessibility[39]. These nucleosome-level effects have functional implications, as CpG-rich exons and CpG islands are known to affect transcription initiation, promoter architecture, and co-transcriptional splicing regulation[40].These findings have highlighted that CpG-rich and CpG-regulated exonic regions are not only sequence-specific features but also structural determinants of chromatin organization, motivating deeper investigation of CpG island overlap within essential gene exons.

Therefore, we examined the regions of CpG islands that overlap with the exons of essential genes encoding essential proteins. These analyses not only provided insight into which portions of the protein-coding regions are associated with CpG islands, but also allowed us to assess the location of enrichment of CpGs-from both CGI and non-CGI regions, across the gene bodies of all human essential genes.

Overall, this study provides a quantitative framework for exploring CpG-driven proteomic architecture and its implications in genomic organization.

## 2. Data acquisition

### 2.1. Protein data

In this study, a total of 3,230 essential proteins from the model organism ‘*homo sapiens*’ were retrieved from the DEG database [41]. Among these proteins, amino acid sequences corresponding to DEG20110914, DEG20110939, and DEG20111255 were neither available in DEG nor in UniProt. In addition, five pairs of entries were identified that shared identical amino acid sequences: DEG20111590 (LCOR) and DEG20111591 (C10orf12); DEG20111741 (RBM14) and DEG20111743 (RBM14); DEG20112241 (CSNK1G1) and DEG20112243 (CSNK1G1); DEG20112422 (TNFSF12) and DEG20112427 (TNFSF12); DEG20113076 (WDR45) and DEG20113081 (WDR45). After accounting for these redundancies and missing sequences, a final dataset of 3,222 unique human essential protein sequences was obtained and used for subsequent analyses.

### 2.2. CpG islands mapping data

#### Obtaining essential genes-associated information

The DEG database was scraped to obtain essential genes-associated UniProt accession numbers and Gene Ref IDs using a Python script. The resulting file was subsequently used to filter the required CCDS IDs and other relevant information from the master BioMart file.

#### Preparation of the master BioMart file

Ensembl BioMart tool[42] was used to prepare the master file. The human genome assembly GRCh38/hg38 was selected as the dataset. The filters (F) and attributes (A) applied for constructing this file were as follows:

F - chromosomes 1–22, X, and Y; protein-coding genes.

A - Chromosome/Scaffold name, ‘nc accession’, ‘gene’, ‘gene id’, ‘ccds id’, ‘ccds status’, ‘cds strand’, ‘cds from’, ‘cds to’, and ‘cds locations’.

The resulting BioMart file contained CCDS IDs for nearly all essential gene transcripts.

#### Intersecting essential gene information with the BioMart file

The list of all 3,230 essential genes was intersected with the BioMart file to retrieve the corresponding CCDS IDs along with their exon locations/coordinates. For genes lacking CCDS IDs in this intersection, their IDs were manually searched in the Ensembl CCDS database, followed by recording the exon coordinates. During this process, a total of 12 genes were removed from the working dataset because some were classified as pseudogenes with incomplete information, while the others had CCDS IDs retracted due to inconsistencies in functional annotation or genomic location. The final curated working database consisted of 3,217 essential genes.

#### Recording exon sequences for essential genes

A set of Python scripts were written to extract exact exon sequences from the fasta files of different chromosomes. Fasta files containing exon sequences with their locations as headers were produced as a result, which were further used for analyses purposes.

#### Overlapping the CGI coordinates with exon sequences

To identify the CGI regions associated with exons for each gene (later defined as the exon overlap fraction), CGI coordinates were mapped onto exon sequences across all genes. For each gene, a highlighted DOCX file was generated in which the nucleotides overlapping CpG island regions within exons were visually marked.

## 3. Methods

### 3.1. CpG-induced amino acids and chromosomal grouping of human essential proteins

To investigate the influence of CpG island composition on amino acid usage and chromosomal distribution, we analyzed 3,222 human essential protein sequences. CpG (Cytosine–phosphate–Guanine) islands were identified within coding regions and promoter-proximal sequences using standard criteria: GC content ≥ 50%, observed/expected CpG ratio ≥ 0.6, and length ≥ 200 bp. Codons containing both cytosine (C) and guanine (G) nucleotides were specifically flagged, as these are more likely to occur within CpG islands. The analysis focused on codons encoding five amino acids—alanine (A), arginine (R), serine (S), proline (P), and threonine (T)—which are frequently induced by CG-containing codons. For each human essential protein, the relative frequency of these five amino acids was calculated, yielding five-dimensional composition vectors. All sequences were then clustered using the k-means clustering algorithm. The optimal number of clusters was determined by maximizing the average *Silhouette* score, which evaluates how well each data point fits within its assigned cluster relative to neighboring clusters [43]. The score ranges from –1 to +1, with higher values indicating more cohesive clustering [44]. Following clustering, each protein was mapped to its corresponding chromosome using gene annotations from Ensembl and UniProt [45, 46]. Human essential proteins were grouped according to their encoding chromosome, resulting in 23 chromosomal clusters (22 autosomes plus chromosomes X) [47]. It was noted that no gene/protein in chromosome Y was found to designated as essential gene/protein as reported [41, 47].

### 3.2. Shannon entropy estimation of human essential proteins

To quantify sequence-level compositional complexity, Shannon entropy was computed for each protein sequence *S*_*j*_ ∈ 𝒮, where 𝒮 denotes the set of all curated human essential proteins. The analysis considered the 20 canonical amino acids:

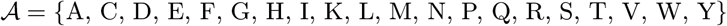

For a sequence of length *L*_*j*_, let 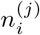 denote the number of occurrences of amino acid *a*_*i*_ ∈ 𝒜 in sequence *S*_*j*_. The frequency of *a*_*i*_ is then defined as:

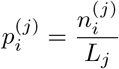

The Shannon entropy *H*_*j*_ for sequence *S*_*j*_ is computed as:

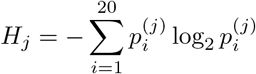

with the convention that 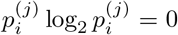 whenever 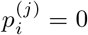. This formulation yields a scalar measure of amino acid diversity per sequence. Entropy values were computed independently for each protein sequence, resulting in a distribution 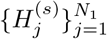 corresponding to a Cluster s containing multiple sequences.

To assess whether entropy distributions differed significantly between two Cluster m and Cluster n, a Wilcoxon rank-sum test was applied under the null hypothesis:

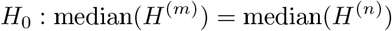

The test was implemented in MATLAB R2025b using the ranksum function. Entropy distributions were visualized using normalized histograms and kernel density estimates (KDE), with Gaussian kernels and bandwidth selection via MATLAB’s default rule. KDE curves and annotated histograms were overlaid to highlight distributional features, including modality, spread, and central tendency. Summary statistics (mean, standard deviation, sample size, and *p*-value) were embedded directly in the plots for interpretability.

### 3.3. CpG-induced amino acid composition and correlation cnalysis

The relative percentage composition of five amino acids—Alanine (A), Arginine (R), Serine (S), Proline (P), and Threonine (T)—from 3222 human essential proteins were computed. Each protein was represented by a five-dimensional vector corresponding to the normalized abundance of these residues. Descriptive statistics, including mean, median, and variance, were computed for each amino acid to assess distributional properties. Outliers were identified using the interquartile range (IQR) method, where values falling below *Q*_1_ − 1.5 × IQR or above *Q*_3_ + 1.5 × IQR were flagged [48]. Pairwise relationships among amino acids were evaluated using Pearson correlation coefficients.

### 3.4. Detection and visualization of CpG-induced amino acid regions in human essential proteins

To identify CpG-influenced regions within protein sequences derived from chromosome-specific datasets, we implemented a sliding window-based approach that mimics CpG island detection at the nucleotide level. Protein sequences were retrieved from curated FASTA files corresponding to individual chromosomes. Each sequence was scanned using a fixed-length window of 67 amino acids, approximating the length of a canonical CpG island ( ∼ 200 bp) [2].

Amino acids encoded by codons containing CpG di-nucleotides n(C and G) – namely arginine (R), threonine (T), proline (P), alanine (A), and serine (S)—were designated as CpG-influenced. For each window, the fraction of CpG-influenced residues was computed. Regions where this fraction exceeded a predefined threshold (≥ 0.5) were classified as CpG-enriched [2].

For each protein, the density of CpG-influenced residues was plotted across the sequence length. Enriched regions were highlighted using shaded overlays, and a horizontal reference line denoted the enrichment threshold [49]. To facilitate downstream analysis, metadata including sequence headers, start and end positions of enriched regions, fractional scores, and amino acid segments were compiled.

This pipeline was applied systematically across all chromosome-specific protein sets, enabling comparative visualization and quantification of CpG-influenced domains at the proteomic level.

#### 3.4.1. CpG metric for quantitative analysis of CpG induced regions in human essential proteins

For each human essential protein, CpG-induced amino acid region-based metrics were computed, including:

##### 1. Average CpG density

Let a protein sequence contain *n* CpG-induced amino acid regions, each denoted by *R*_*i*_ for *i* = 1, 2, …, *n*. A CpG-induced region is defined as a contiguous subsequence enriched in CpG-associated amino acids:

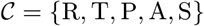

where R = Arginine, T = Threonine, P = Proline, A = Alanine, and S = Serine.

For each region *R*_*i*_ of length *L*_*i*_ (in amino acids), let *C*_*i*_ be the count of residues in *R*_*i*_ that belong to the CpG-associated set 𝒞. Then, the CpG density of region *R*_*i*_ is defined as:

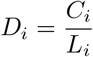

The average CpG density for the protein is given by:

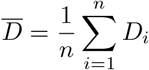

##### 2. Length of the longest CpG-induced amino acid region

Let *L*_*i*_ denote the length of the *i*^*th*^ CpG-induced region. Then, the longest region length is:

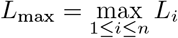

##### 3. Fraction of the amino acid sequence covered

Let *L*_total_ be the total length of the protein sequence. The fraction of the sequence covered by CpG-induced regions is:

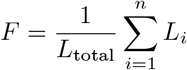

For each cluster, all the three aforementioned metrics were aggregated chromosome-wise. Descriptive statistics including mean, median, and standard deviation were computed for each metric per chromosome. These summaries were used to assess intra-cluster variability and inter-chromosomal trends. Separate histogram plots for each metric were created for Cluster-1 and Cluster-2 to visualize distributional differences.

For each human essential protein, CpG-induced amino acid region-based metrics were computed and aggregated chromosome-wise within each cluster. The following three grouped metrics were defined:

##### 1. Average CpG Density (Mean and Standard Deviation)

Let a protein *P*_*j*_ contain *n*_*j*_ CpG-induced regions *R*_*jl*_, each with CpG density 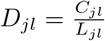, where *C*_*jl*_ is the count of CpG-associated residues and *L*_*jl*_ is the region length. The average CpG density for *P*_*j*_ is:

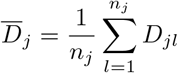

For chromosome *i* and cluster *k*, with *n*_*ik*_ proteins, the chromosome-wise mean and standard deviation are:

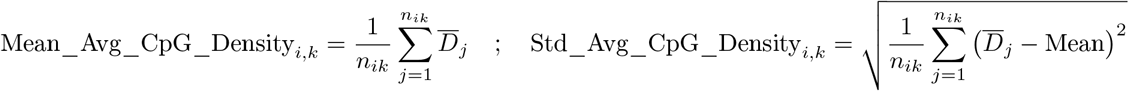

##### 2. Length of the Longest CpG-Induced Region (Mean and Standard Deviation)

Let 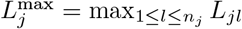 be the longest CpG-induced region in protein *P*_*j*_. Then:

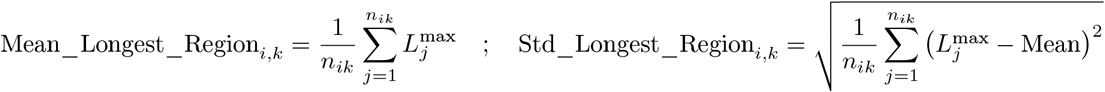

##### 3. Fraction of Protein Sequence Covered (Mean and Standard Deviation)

Let 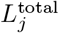 be the total length of protein *P*_*j*_, and 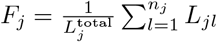 be the fraction covered by CpG-induced regions. Then:

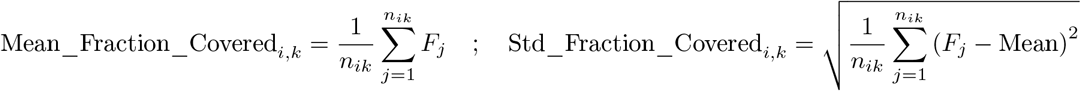

These metrics were used to quantify CpG motif enrichment, segment continuity, and coverage across chromosomes and clusters. Histogram plots and line charts were generated to visualize distributional differences.

#### 3.4.2. Motif Composition Quantification of the Longest CpG-Influenced Amino Acid Regions

Among the CpG-induced regions identified for human essential proteins across chromosomes and clusters, the longest segment per protein was selected based on maximal length. This ensured a consistent and non-redundant representation of CpG-enriched content for downstream motif analysis.

Each longest CpG-induced region was analyzed for amino acid composition using three motif metrics:

- **PolarRatio**: Defined as the ratio of the number of polar residues to non-polar residues within the longest CpG-induced region. Polar residues include Serine (S), Threonine (T), Asparagine (N), Glutamine (Q), Tyrosine (Y), Cysteine (C), Aspartic acid (D), Glutamic acid (E), Histidine (H), Lysine (K), and Arginine (R). Non-polar residues include Alanine (A), Valine (V), Leucine (L), Isoleucine (I), Methionine (M), Phenylalanine (F), Tryptophan (W), Proline (P), and Glycine (G).
- **AcidicFrac**: Fraction of acidic residues — Aspartic acid (D) and Glutamic acid (E) — relative to the total length of the CpG-induced region.
- **BasicFrac**: Fraction of basic residues — Arginine (R), Lysine (K), and Histidine (H) — relative to the total length of the CpG-induced region.

For each human essential protein, these metrics were computed for its longest CpG-influenced region and aggregated chromosome-wise to assess motif distribution patterns. For each cluster, chromosome-wise histograms were generated for PolarRatio, AcidicFrac, and BasicFrac to capture intra-chromosomal variability.

To isolate proteins exhibiting pronounced motif enrichment or depletion, threshold-based filtering was applied to the computed metrics. Proteins were selected if they satisfied any of the following criteria:

- **Polar-rich**: PolarRatio ≥ 0.8
- **Polar-depleted**: PolarRatio ≤ 0.2
- **Acidic-rich**: AcidicFrac ≥ 0.2
- **Basic-rich**: BasicFrac ≥ 0.2

These thresholds were empirically chosen to highlight proteins with extreme motif biases, facilitating targeted annotation and comparative analysis. Filtered subsets were stratified by chromosome and cluster.

Motif composition trends and distribution patterns were visualized to enable comparative analysis across chromosomes and between clusters. For each cluster, line plots were generated to depict the mean motif fractions — PolarRatio, AcidicFrac, and BasicFrac — across chromosomes 1 through 22 and X, revealing global trends and cluster-specific biases. Heatmaps were constructed to represent chromosome-wise motif densities for polar, acidic, and basic residues, using color gradients to highlight enrichment patterns and inter-chromosomal variability. Additionally, histograms were produced for each cluster to illustrate the distribution of PolarRatio, AcidicFrac, and BasicFrac values across individual chromosomes, capturing intra-chromosomal variation in motif enrichment.

### 3.5. Cluster-wise subgroup identification of human essential proteins

Essential proteins from each human chromosome were analyzed using three CpG-associated metrics: CpG density, fraction of the sequence covered by CpG islands, and the length of the longest uninterrupted CpG-induced amino acid region. Prior to clustering, all three metrics were standardized using z-score normalization to ensure equal weighting and comparability across chromosomes.

K-means algorithm was applied on the standardized CpG metric associated to each chromosome-specific set of human essential proteins to obtain further subgroups. The optimal number of clusters (*k*) was determined by evaluating average silhouette scores across *k* = 2 to *k* = 10; in all cases, *k* = 2 yielded the highest silhouette score, confirming a stable bipartite structure of each chromosome in each cluster. Subgroup assignments were recorded per chromosome and used for comparative analysis of CpG metric variability and structural stratification.

### 3.6. CpG islands mapping and CG distribution in human essential genes

#### 3.6.1. Calculation of exon overlap fraction

The exon overlap fraction was computed by dividing the total number of highlighted nucleotides across all exons of a gene by the total nucleotide count of all exons in that gene:

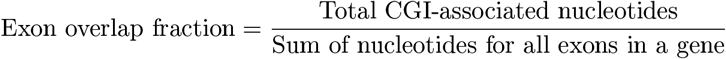

A histogram of exon overlap fractions was generated using these values. Next, CGI-associated CpG counts were calculated from the CGI-overlapping stretches of exons, while the remaining exon nucleotides were considered non–CGI-associated regions; CpGs present in these segments were defined as non-CGI-associated CpGs. All genes were then distributed into 10 bins based on their exon overlap fraction values, and for each bin, normalized CGI-associated and non-CGI-associated CpG counts were calculated as follows:

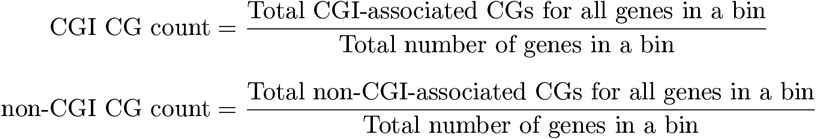

#### 3.6.2. CpG density analyses across human essential genes

Prior to calculating CpG density across genes, strand information was retrieved for all genes to determine their genomic orientation. Once strand orientation was established, CpG sites were scored in the concatenated exon sequences using a simple algorithm. Genes located on the minus strand were flipped before scoring. All CG dinucleotides were assigned a score of “1-1,” whereas non-CG sites were assigned a score of 0. This procedure was applied uniformly across all essential gene–associated exon sequences.

Following scoring, CGI-associated and non-CGI-associated CpG densities were plotted in the 5^*′*^ → 3^*′*^ direction by scaling all genes to an identical length corresponding to the smallest gene.

The same plotting strategy was also applied to Classes 1 and 2, where Class-1 represents bins with exon-overlap-fractions ranging between 0-0.5, while, Class-2 covers fractions from 0.5 to 1.

## 4. Results and analyses

### 4.1. Hierarchical grouping of human essential proteins by CpG-Induced amino acid composition and chromosomal origin

A total of 3,222 human essential proteins were first clustered into two groups using the k-means algorithm, based on the relative frequencies of five CpG-associated amino acids: alanine (A), arginine (R), serine (S), proline (P), and threonine (T). The optimal number of clusters (k = 2) was selected by maximizing the average *Silhouette* score (0.46), indicating moderately distinct separation.

Cluster 1 comprised 991 proteins, while Cluster 2 contained the remaining 2,231 sequences (**Supplementary file-1, and Supplementary file-2**). To further explore the structure of compositional variability, we performed Principal Component Analysis (PCA) on the relative percentage composition of five CpG-associated amino acids—Alanine (A), Arginine (R), Serine (S), Proline (P), and Threonine (T)—across 3,222 human essential proteins. Each protein was represented as a five-dimensional vector, capturing its CpG-driven amino acid profile.

The first two principal components explained a substantial proportion of the total variance: PC1 accounted for 38.98% and PC2 for 22.64%. PC1 was primarily influenced by the inverse relationship between Serine and Alanine, while PC2 reflected opposing contributions from Proline and Threonine. These axes captured dominant patterns of amino acid co-variation, consistent with the correlation structure reported earlier (Figure 6).

The PCA scatter plot (Figure 1) revealed two distinct compositional regimes. Red circles and blue crosses represent proteins assigned to Cluster 1 and Cluster 2, respectively, based on k-means clustering of CpG-associated amino acid frequencies. The clear separation between clusters in PCA space supports the hypothesis that CpG-driven amino acid usage is structured and non-random, shaped by genomic context and functional constraints.

**Figure 1:**
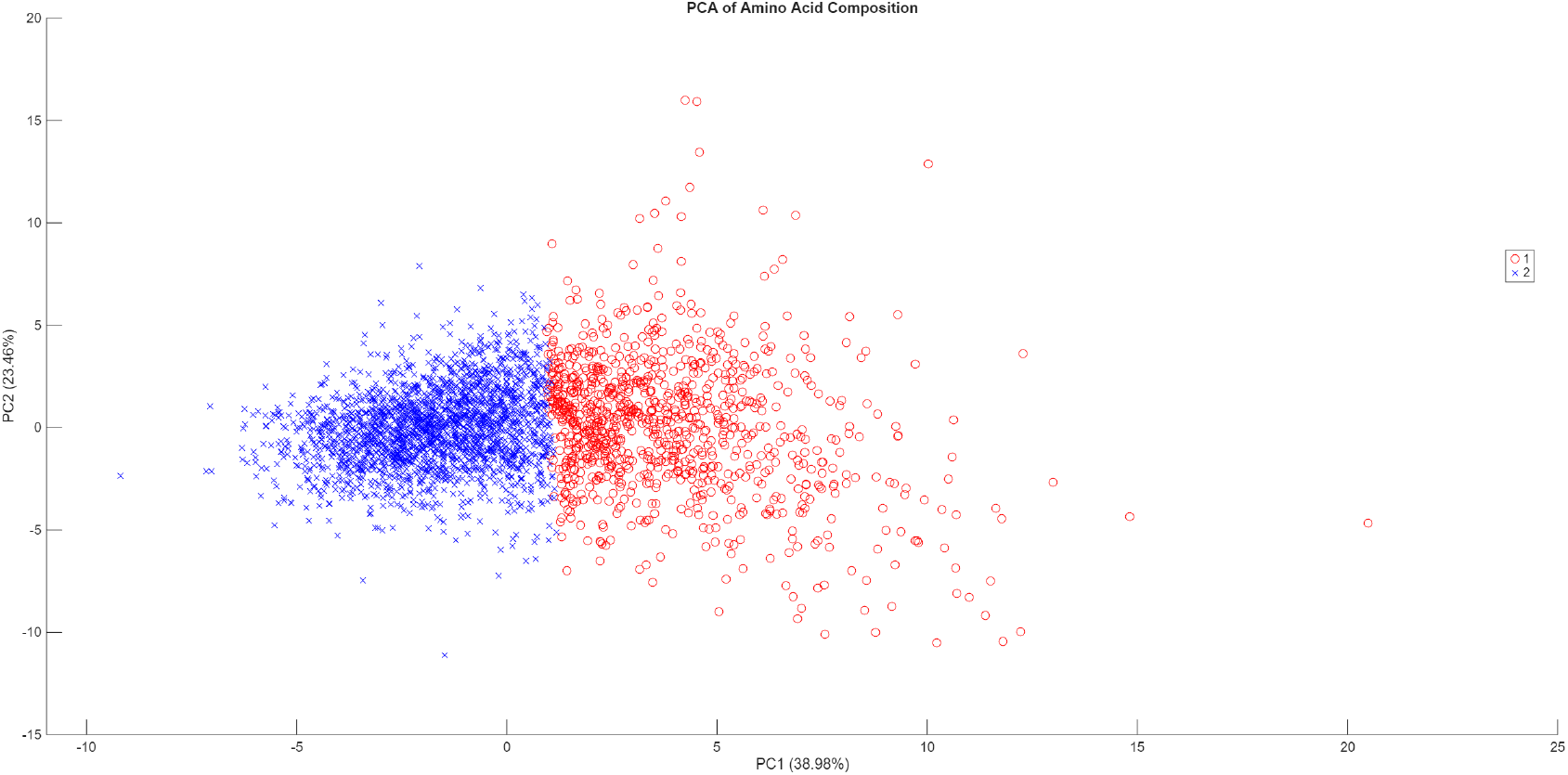
Principal component analysis of CpG-associated amino acid composition. PCA scatter plot of 3,222 human essential proteins based on relative frequencies of five CpG-associated amino acids (A, R, S, P, T). PC1 and PC2 explain 38.98% and 22.64% of the total variance, respectively. Red circles and blue crosses denote proteins assigned to Cluster 1 and Cluster 2 via k-means clustering. The compositional separation along PC1 and PC2 reflects underlying amino acid trade-offs and supports the presence of distinct CpG-driven regimes.

Each cluster was subsequently subdivided according to chromosomal origin, using gene annotations. This resulted in 23 chromosome-based subgroups within each cluster (22 autosomes plus chromosome X), as no essential protein sequences were mapped to chromosome Y (**Supplementary file-1, and Supplementary file-2**). This hierarchical organization enabled comparative analysis of CpG-induced amino acid regions across both compositional clusters and chromosomal contexts, revealing potential links between genomic architecture and essential protein function.

Analysis of absolute and relative frequencies revealed distinct chromosomal biases between clusters (Table 1). Chromosome 1 contributed the highest number of proteins overall (320), with 104 in Cluster 1 and 216 in Cluster 2, corresponding to 10.49% and 9.68% of their respective totals. Chromosomes 19 and 17 showed elevated representation in Cluster 1 (7.16% and 7.06%), suggesting enrichment of CpG-associated amino acid usage. In contrast, chromosome X exhibited a bias toward Cluster 2, contributing 7.84% of its proteins compared to 5.45% in Cluster 1. Chromosomes 4, 14, and 22 also showed higher relative frequencies in Cluster 2, indicating potential compositional divergence (Figure 2).

**Table 1:** Chromosome-wise distribution of human essential proteins across CpG-based clusters.

| Chromosome | Cluster 1 (Count) | Cluster 2 (Count) | Cluster 1 (%) | Cluster 2 (%) |
| --- | --- | --- | --- | --- |
| Chromosome 1 | 104 | 216 | 10.49 | 9.68 |
| Chromosome 2 | 56 | 154 | 5.65 | 6.90 |
| Chromosome 3 | 51 | 140 | 5.15 | 6.28 |
| Chromosome 4 | 24 | 83 | 2.42 | 3.72 |
| Chromosome 5 | 51 | 115 | 5.15 | 5.16 |
| Chromosome 6 | 41 | 101 | 4.14 | 4.53 |
| Chromosome 7 | 45 | 99 | 4.54 | 4.44 |
| Chromosome 8 | 27 | 67 | 2.73 | 3.00 |
| Chromosome 9 | 40 | 92 | 4.04 | 4.12 |
| Chromosome 10 | 35 | 87 | 3.53 | 3.90 |
| Chromosome 11 | 61 | 124 | 6.16 | 5.56 |
| Chromosome 12 | 67 | 129 | 6.76 | 5.78 |
| Chromosome 13 | 15 | 44 | 1.51 | 1.97 |
| Chromosome 14 | 25 | 83 | 2.52 | 3.72 |
| Chromosome 15 | 29 | 75 | 2.93 | 3.36 |
| Chromosome 16 | 38 | 71 | 3.84 | 3.18 |
| Chromosome 17 | 70 | 124 | 7.06 | 5.56 |
| Chromosome 18 | 20 | 38 | 2.02 | 1.70 |
| Chromosome 19 | 71 | 96 | 7.16 | 4.30 |
| Chromosome 20 | 33 | 59 | 3.33 | 2.65 |
| Chromosome 21 | 8 | 19 | 0.81 | 0.85 |
| Chromosome 22 | 26 | 40 | 2.62 | 1.79 |
| Chromosome X | 54 | 175 | 5.45 | 7.84 |
| **Total** | **991** | **2231** | **100.00** | **100.00** |

**Figure 2:**
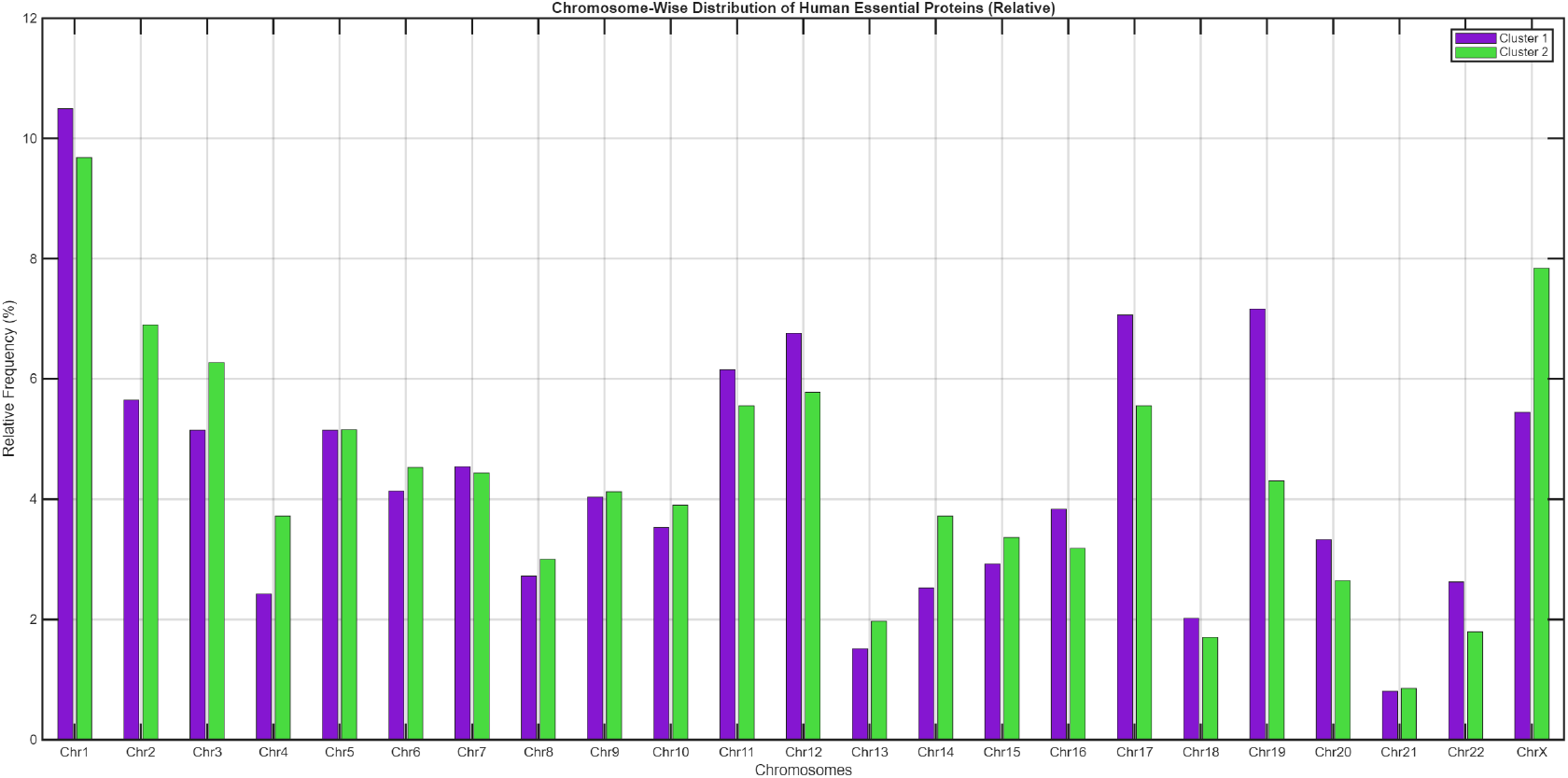
Chromosome-wise distribution of human essential proteins (relative frequencies). Grouped bar chart showing the relative frequency (%) of essential proteins per chromosome within each CpG-based cluster. Cluster 1 and Cluster 2 were derived from k-means clustering based on the frequency of CpG-associated amino acids (A, R, S, P, T). Chromosomes 19 and 17 exhibit enrichment in Cluster 1, while chromosome X shows a pronounced bias toward Cluster 2.

### 4.2. Cluster-wise Shannon entropy of human essential proteins

To evaluate the compositional diversity of human essential proteins, Shannon entropy was computed for each sequence and compared across the two CpG-driven clusters, viz. Cluster 1 and Cluster 2. The resulting entropy distributions revealed a statistically significant shift in sequence complexity between clusters (Figure 3).

**Figure 3:**
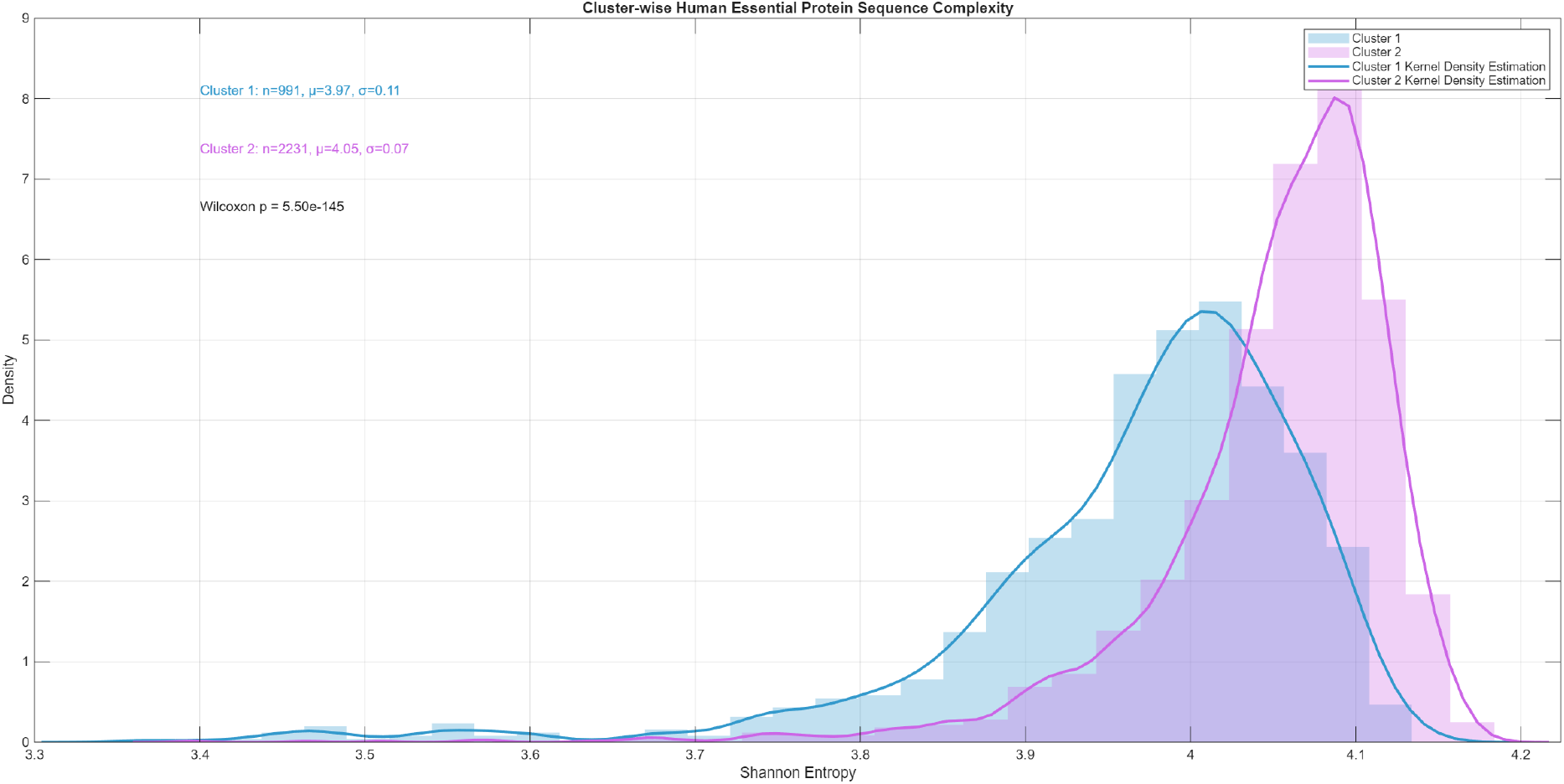
Density plot showing the distribution of Shannon entropy values for 3,222 human essential protein sequences, partitioned into two clusters based on CpG-associated amino acid composition. Cluster 1 (blue) and Cluster 2 (pink) are visualized using overlaid normalized histograms and kernel density estimates (KDE). Annotated statistics indicate sample size (*n*), mean entropy (*µ*), and standard deviation (*σ*) for each cluster. Cluster 1: *n* = 991, *µ* = 3.91, *σ* = 0.11; Cluster 2: *n* = 2231, *µ* = 4.05, *σ* = 0.07. A Wilcoxon rank-sum test revealed a highly significant difference in entropy distributions (*p* = 5.50 *×* 10^−145^), suggesting distinct compositional regimes between clusters.

Cluster 1 proteins exhibited lower entropy values overall, with a mean of *µ*_1_ = 3.91 and standard deviation *σ*_1_ = 0.11 across *n*_1_ = 961 sequences. In contrast, Cluster 2 proteins showed higher compositional diversity, with a mean of *µ*_2_ = 4.05 and standard deviation *σ*_2_ = 0.07 across *n*_2_ = 2253 sequences. The difference in distributions was highly significant, as assessed by a Wilcoxon rank-sum test (*p* = 5. 50 × 10^−145^), rejecting the null hypothesis of equal medians.

Kernel density estimation (KDE) and overlaid histograms further illustrated the separation in entropy profiles. Cluster 1 displayed a broader, left-skewed distribution, while Cluster 2 was more sharply peaked around its higher mean. These results suggest that CpG-associated amino acid usage not only partitions proteins into distinct compositional regimes, but also reflects underlying constraints on sequence variability. The observed entropy shift may be indicative of differential functional constraints, structural preferences, or evolutionary pressures acting on the two human essential protein subsets.

### 4.3. CpG-associated amino acid composition and distributional analysis

Relative percentage composition of five CpG-associated amino acids—Alanine (A), Arginine (R), Serine (S), Proline (P), and Threonine (T)—was analyzed across 3,222 human essential proteins (cluster-wise) to investigate compositional biases and inter-residue relationships.

In Cluster 1 (*n* = 991 human essential proteins), Serine and Proline exhibited markedly elevated mean compositions (10.46% and 9.65%, respectively), with Serine also showing the highest median (10.34%) and Proline the highest variance (8.69), indicating substantial heterogeneity across proteins (Table 2). Alanine and Arginine maintained moderate means (7.38% and 5.87%) with variances exceeding 5.0, while Threonine displayed the lowest variance (2.61), suggesting a more conserved distribution. Outlier analysis based on the interquartile range (IQR) method identified Proline (*n* = 59) and Arginine (*n* = 44) as the most frequent outliers, reflecting context-specific enrichment in select protein subsets. Alanine and Serine each showed 25 outliers, and Threonine had 27, consistent with their relatively stable distributions within the cluster.

**Table 2:** Descriptive statistics and outlier counts for Cluster 1 (*n* = 991)

| Amino Acid | Mean (%) | Median (%) | Standard deviation | Outlier Count |
| --- | --- | --- | --- | --- |
| Alanine (A) | 7.38 | 7.09 | 2.24 | 25 |
| Arginine (R) | 5.90 | 5.51 | 2.42 | 44 |
| Serine (S) | 10.46 | 10.34 | 2.74 | 25 |
| Proline (P) | 9.68 | 9.03 | 2.95 | 59 |
| Threonine (T) | 5.36 | 5.23 | 1.61 | 27 |

Cluster 2 (*n* = 2, 231 human essential proteins) demonstrated lower overall mean compositions and reduced variability for all five CpG-associated amino acids (Table 3). Serine remained the most abundant (mean = 7.44%), followed by Alanine (6.65%) and Arginine (5.69%), while Proline showed the lowest mean (5.04%) and variance (2.19), indicating a tightly regulated distribution. Threonine exhibited the lowest variance overall (1.55), reinforcing its compositional stability. Outlier counts were highest for Arginine (*n* = 77) and Alanine (*n* = 60), suggesting selective enrichment in specific protein contexts. Serine and Threonine showed moderate outlier frequencies (24 and 49, respectively), while Proline had the fewest (*n* = 16), consistent with its constrained variability across the cluster.

**Table 3:** Descriptive statistics and outlier counts for Cluster 2 (*n* = 2, 231)

| Amino Acid | Mean (%) | Median (%) | Standard deviation | Outlier Count |
| --- | --- | --- | --- | --- |
| Alanine (A) | 6.65 | 6.43 | 1.91 | 60 |
| Arginine (R) | 5.69 | 5.40 | 1.87 | 77 |
| Serine (S) | 7.44 | 7.33 | 1.91 | 24 |
| Proline (P) | 5.04 | 5.04 | 1.48 | 16 |
| Threonine (T) | 5.21 | 5.19 | 1.24 | 49 |

Histogram-based visualization of relative percentage compositions for five CpG-associated amino acids—Alanine (A), Arginine (R), Serine (S), Proline (P), and Threonine (T)—revealed distinct distributional patterns between Cluster 1 (*n* = 991) and Cluster 2 (*n* = 2, 231) human essential proteins (Figure 4 and Figure 5).

**Figure 4:**
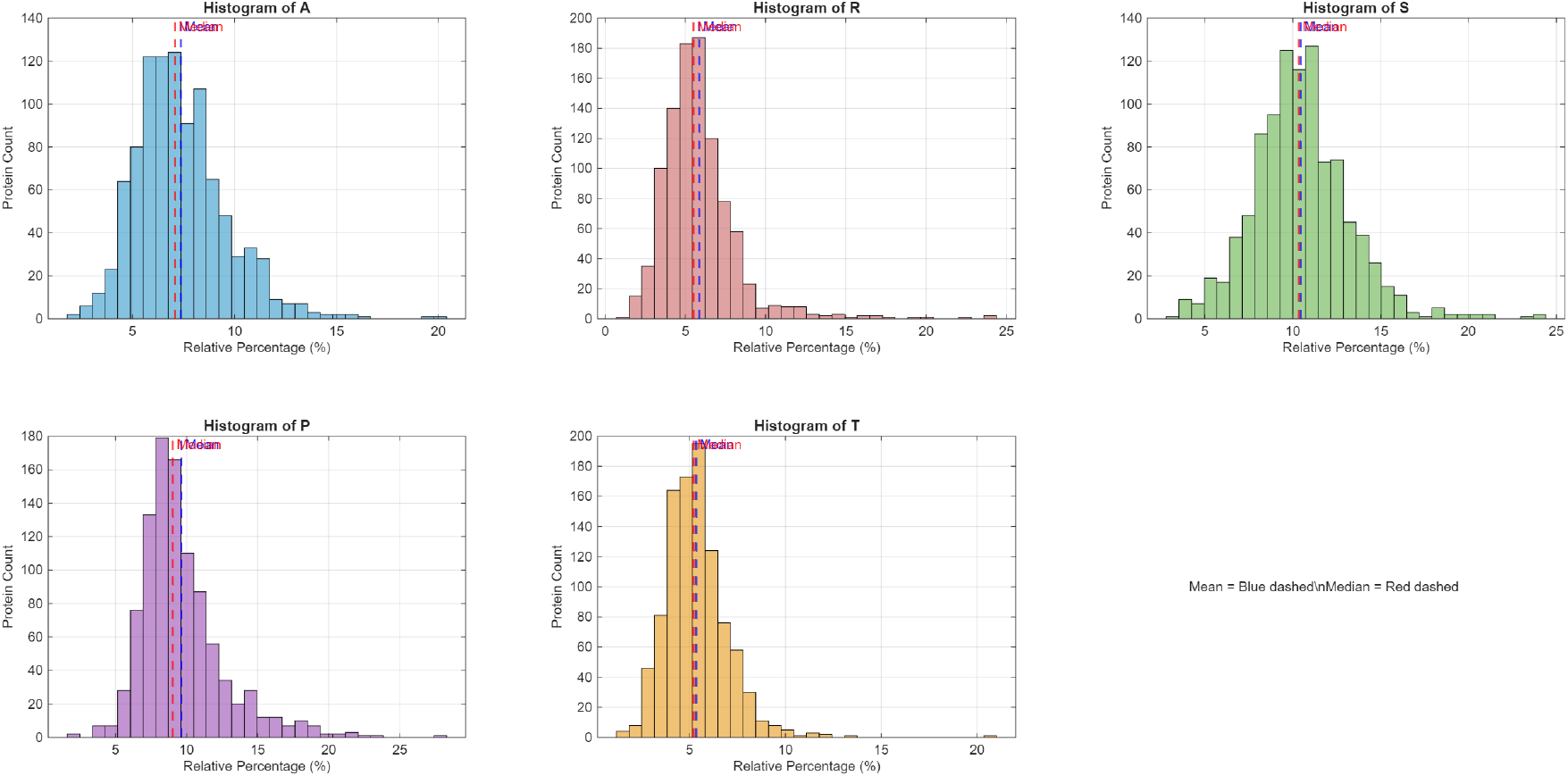
Histograms of relative percentage compositions for five CpG-associated amino acids in Cluster 1 (*n* = 991). Dashed vertical lines indicate mean values for each group.

**Figure 5:**
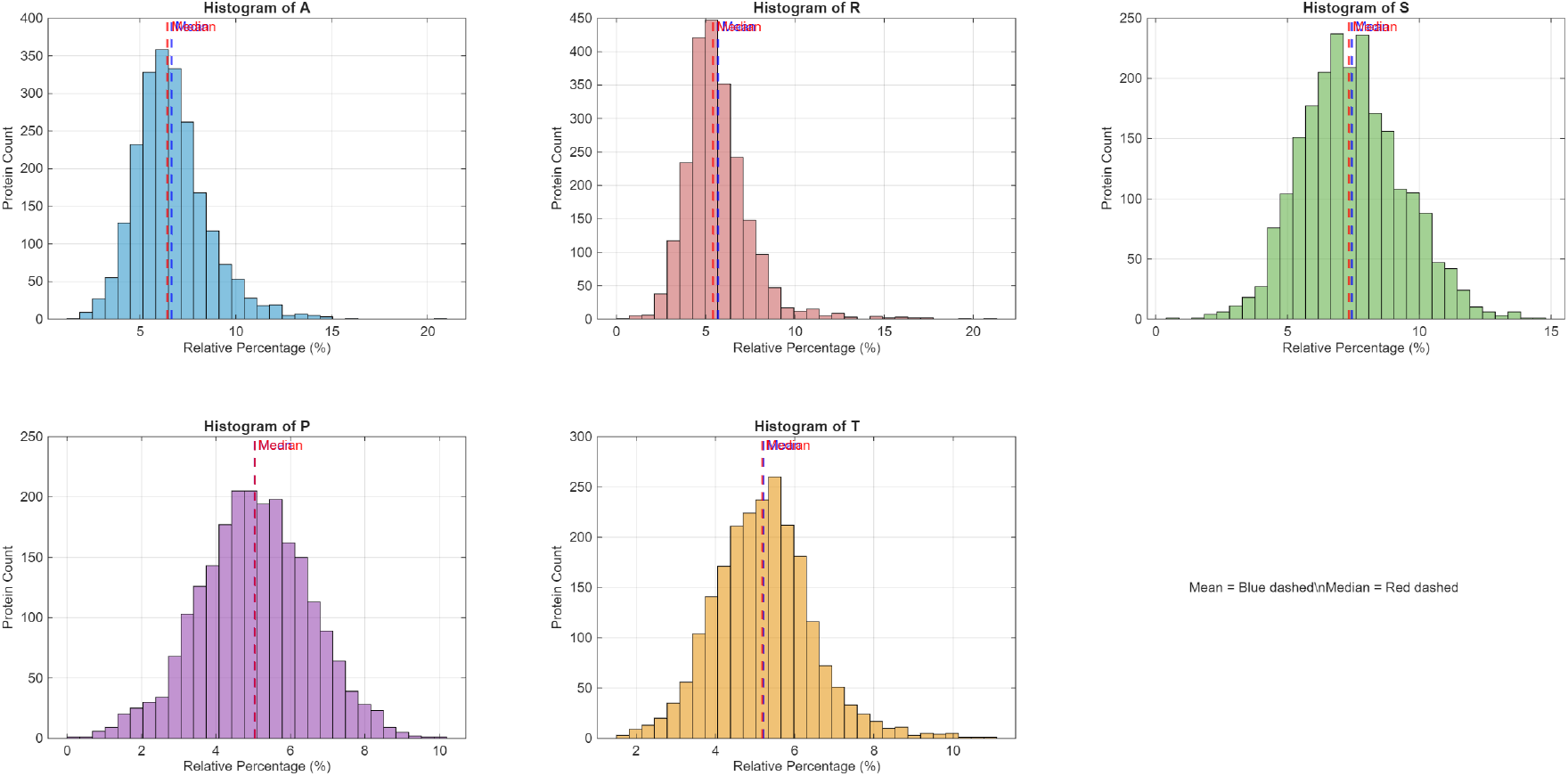
Histograms of relative percentage compositions for five CpG-associated amino acids in Cluster 2 (*n* = 2, 231). Dashed vertical lines indicate mean values for each group.

**Figure 6:**
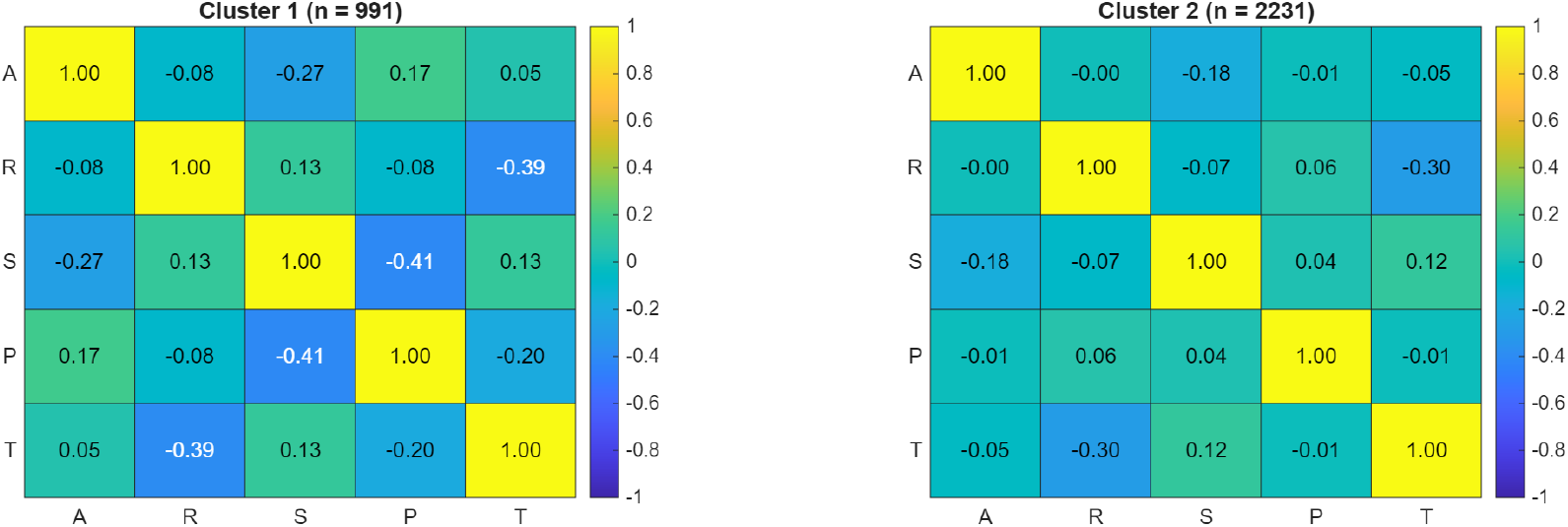
Heatmaps of Pearson correlation matrices for CpG-associated amino acid compositions in Cluster 1 (*n* = 991) and Cluster 2 (*n* = 2, 231). Color intensity reflects correlation strength; blue indicates negative correlation, yellow indicates positive correlation.

Cluster 1 exhibited broader and right-shifted distributions for Serine and Proline, with visibly higher mean values and wider spread, consistent with elevated variance and outlier counts. The frequency peaks for Serine and Proline were more dispersed, indicating compositional heterogeneity across proteins. Alanine also showed a moderately right-shifted distribution, while Threonine and Arginine maintained narrower profiles with moderate spread. Mean values, indicated by dashed vertical lines, were consistently higher in Cluster 1 for Serine, Proline, and Alanine.

In contrast, Cluster 2 displayed narrower distributions with sharper peaks for all five residues, particularly Proline and Threonine, reflecting tighter compositional control and reduced variability. The distributions were more symmetric and left-shifted, with lower mean values across all amino acids. Arginine and Alanine showed the highest outlier counts in Cluster 2, yet their distributions remained more compact compared to Cluster 1. These trends align with the descriptive statistics and outlier profiles, reinforcing cluster-specific compositional biases and suggesting functional or structural divergence among protein subsets.

Inter-residue dependencies were assessed using Pearson correlation matrices derived from the relative percentage compositions of five CpG-associated amino acids—Alanine (A), Arginine (R), Serine (S), Proline (P), and Threonine (T)—across Cluster 1 (*n* = 991) and Cluster 2 (*n* = 2, 231) human essential proteins. Figure 6 visualizes the resulting correlation structures, revealing distinct compositional dynamics between the two clusters.

In Cluster 1, several moderate correlations were observed, including a negative association between Serine and Proline (*r* = − 0.41) and between Arginine and Threonine (*r* = − 0.39), suggesting mutually exclusive enrichment patterns (Figure 6). Alanine showed a weak negative correlation with Serine (*r* = − 0.27) and a modest positive correlation with Proline (*r* = 0.17). These patterns reflect a more heterogeneous and interdependent compositional landscape, consistent with the broader distributions and higher outlier frequencies observed in Cluster 1.

In contrast, Cluster 2 exhibited generally weaker correlations, with most pairwise coefficients near zero. The strongest relationship was a moderate negative correlation between Arginine and Threonine (*r* = − 0.30), while Serine showed weak positive associations with Proline (*r* = 0.04) and Threonine (*r* = 0.12) (Figure 6). Alanine displayed minimal correlation with all other residues, indicating a more compositionally constrained and independent distribution. The reduced inter-residue coupling in Cluster 2 aligns with its narrower histograms and lower variance profiles.

### 4.4. CpG island induced regions in human essential proteins

A total of 3222 human essential proteins were analyzed, comprising 1524 CpG-depleted and 1698 CpG-induced proteins. Among the CpG-depleted proteins, 75 (4.92%) were assigned to Cluster-1 and 1449 (95.08%) to Cluster-2. In contrast, the CpG-induced proteins included 916 (53.95%) in Cluster-1 and 782 (46.05%) in Cluster-2 (Table 4). Cluster-2 proteins were consistently abundant across all chromosomes, with Chromosome-01, Chromosome-02, and Chromosome-X contributing the highest counts. In contrast, Cluster-1 proteins were sparsely represented, with no entries detected on Chromosome-14 and Chromosome-22. The complete per-chromosome breakdown is provided in Table 4. It was noteworthy that across all chromosomes, a strictly monotonic pattern was observed in the distribution of CpG-associated proteins: Cluster 1 consistently exhibited a higher frequency of CpG-induced proteins compared to CpG-depleted ones, whereas Cluster 2 showed the opposite trend, with CpG-depleted proteins predominating. This trend was quantitatively supported by the ratio of CpG-depleted to CpG-induced protein frequencies, which remained less than 1 for Cluster 1 and greater than 1 for Cluster 2 across all chromosomes (Table 4). This asymmetry suggests a functional bifurcation in CpG-driven protein architecture, wherein Cluster 1 may be enriched for transcriptionally active or epigenetically responsive proteins, while Cluster 2 may reflect structural or constitutively expressed proteins with reduced CpG content. Furthermore, these findings suggest that CpG depletion among essential proteins is not uniformly distributed across the genome, with certain chromosomes—particularly Chromosome-X—showing elevated insulation from CpG-mediated modulation.

**Table 4:** Distribution of CpG-depleted and CpG-induced proteins across chromosomes and clusters.

| Chromosome | Cluster 1 |  |  | Cluster 2 |  |  |
| --- | --- | --- | --- | --- | --- | --- |
|  | Frequency of CpG-depleted | Frequency of CpG-induced | Ratio of the frequency of CpG Depleted and CpG induced | Frequency of CpG-depleted | Frequency of CpG-induced | Ratio of the frequency of CpG Depleted and CpG induced |
| Chromosome 01 | 8 | 96 | 0.083 | 140 | 76 | 1.842 |
| Chromosome 02 | 3 | 53 | 0.057 | 95 | 59 | 1.610 |
| Chromosome 03 | 5 | 46 | 0.109 | 91 | 49 | 1.857 |
| Chromosome 04 | 2 | 22 | 0.091 | 55 | 28 | 1.964 |
| Chromosome 05 | 5 | 46 | 0.109 | 81 | 34 | 2.382 |
| Chromosome 06 | 6 | 35 | 0.171 | 58 | 43 | 1.349 |
| Chromosome 07 | 2 | 43 | 0.047 | 67 | 32 | 2.094 |
| Chromosome 08 | 4 | 23 | 0.174 | 40 | 27 | 1.481 |
| Chromosome 09 | 1 | 39 | 0.026 | 64 | 28 | 2.286 |
| Chromosome 10 | 2 | 33 | 0.061 | 61 | 26 | 2.346 |
| Chromosome 11 | 4 | 57 | 0.070 | 80 | 44 | 1.818 |
| Chromosome 12 | 6 | 61 | 0.098 | 89 | 40 | 2.225 |
| Chromosome 13 | 2 | 13 | 0.154 | 29 | 15 | 1.933 |
| Chromosome 14 | 0 | 25 | 0.000 | 51 | 32 | 1.594 |
| Chromosome 15 | 4 | 25 | 0.160 | 49 | 26 | 1.885 |
| Chromosome 16 | 2 | 36 | 0.056 | 42 | 29 | 1.448 |
| Chromosome 17 | 4 | 66 | 0.061 | 83 | 41 | 2.024 |
| Chromosome 18 | 3 | 17 | 0.176 | 25 | 13 | 1.923 |
| Chromosome 19 | 3 | 68 | 0.044 | 56 | 40 | 1.400 |
| Chromosome 20 | 2 | 31 | 0.065 | 35 | 24 | 1.458 |
| Chromosome 21 | 2 | 6 | 0.333 | 10 | 9 | 1.111 |
| Chromosome 22 | 0 | 26 | 0.000 | 25 | 15 | 1.667 |
| Chromosome X | 5 | 49 | 0.102 | 123 | 52 | 2.365 |

Representative CpG-induced amino acid density profiles for two human essential proteins—*DEG20113097*|*IMAG2D*| and *DEG2011132*|*RLM*| —are shown in Figure 7. These plots highlight regions exceeding the CpG-AA density threshold, indicating localized enrichment of CpG-influenced residues. Complete CpG-AA density plots for all remaining essential proteins are provided in **Supplementary File-3**.

**Figure 7:**
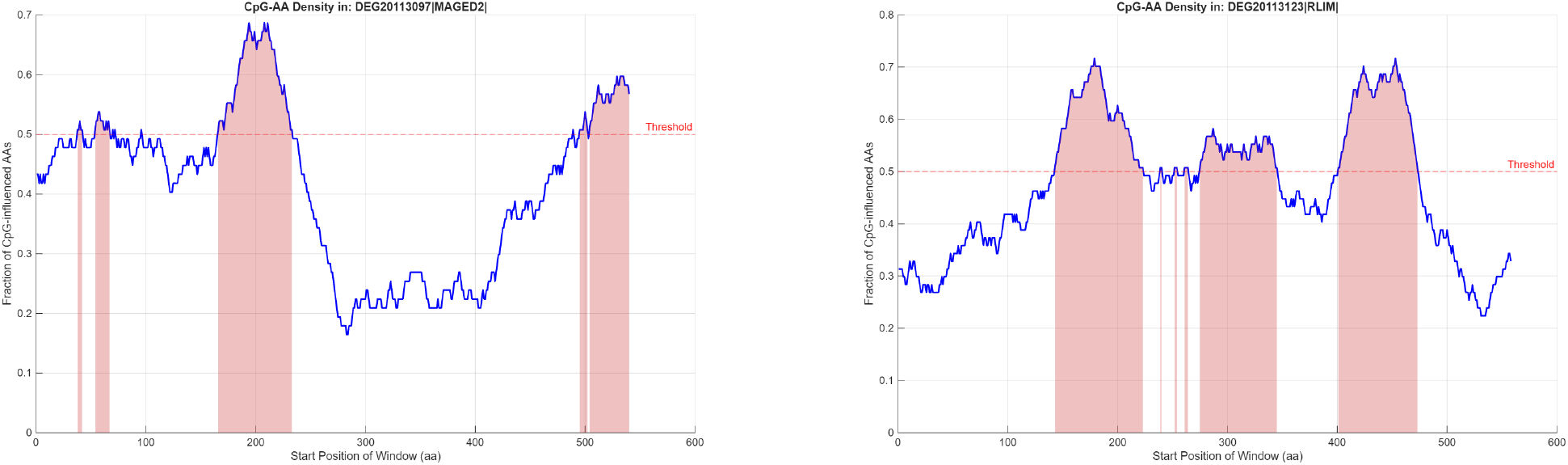
CpG-induced amino acid density profiles for two representative human essential proteins from the chromosome-X: *DEG20113097*|*IMAG2D*| (left) and *DEG2011132*|*RLM* (right). Red-shaded regions indicate windows, where the fraction of CpG-influenced amino acids exceeds the defined threshold.

#### 4.4.1. Chromosome-wise distribution of CpG metrics in Cluster-1 and Cluster-2 human essential proteins

To characterize the structural CpG features of Cluster-1 human essential proteins, we analyzed three key metrics across chromosomes: *average CpG density, length of the longest CpG-induced amino acid region*, and *fraction of sequence covered by CpG-induced amino acid regions*. Histogram plots were generated for each metric, stratified by chromosome (01–22, X), and supported by descriptive statistics (Figures 8, 9, 10 and Table 5). The distribution of average CpG density varied across chromosomes in Cluster-1, with the highest mean observed in Chromosome 3 (0.676) and the lowest in Chromosomes 10 and 21 (0.618) (Table 5). Elevated median values ( ≥ 0.661) were noted in Chromosomes 3, 11, 15, 22, and 23, indicating consistent CpG enrichment among Cluster-1 proteins (Figure 8). Chromosome 8 exhibited the lowest median (0.581), suggesting a subset of proteins having CpG-induced regions with reduced CpG density. The longest CpG-induced amino acid regions were most prominent in Chromosome 16 (mean: 113.08), followed by Chromosomes 19 (97.72) and 13 (96.46), with Chromosome 16 also showing the highest variability (standard deviation: 107.83) (Figure 9, Table 5). Fractional coverage was highest in Chromosomes 16 (mean: 0.203) and 13 (mean: 0.194), while Chromosomes 3, 4, 8, and 10 exhibited lower coverage (mean *<* 0.13) (Figure 10, Table 5). The histograms reveal chromosome-specific patterns in CpG architecture within Cluster-1, with Chromosomes 11, 13, and 16 consistently exhibiting elevated CpG metrics across all three dimensions (Figures 8, 9, 10).

**Table 5:** Chromosome-wise comparison of CpG-induced regions metrics (mean and standard deviation) between Cluster-1 and Cluster-2.

| Chromosome | Mean Avg CpG Density |  | Std Avg CpG Density |  | Mean Longest Region |  | Std Longest Region |  | Mean Fraction Covered |  | Std Fraction Covered |  |
| --- | --- | --- | --- | --- | --- | --- | --- | --- | --- | --- | --- | --- |
|  | Cluster-1 | Cluster-2 | Cluster-1 | Cluster-2 | Cluster-1 | Cluster-2 | Cluster-1 | Cluster-2 | Cluster-1 | Cluster-2 | Cluster-1 | Cluster-2 |
| 1 | 0.639 | 0.622 | 0.119 | 0.162 | 79.969 | 41.289 | 67.599 | 57.181 | 0.168 | 0.055 | 0.119 | 0.055 |
| 2 | 0.637 | 0.627 | 0.119 | 0.175 | 88.151 | 34.203 | 78.739 | 42.382 | 0.169 | 0.040 | 0.140 | 0.041 |
| 3 | 0.676 | 0.642 | 0.144 | 0.175 | 68.587 | 30.163 | 67.756 | 30.882 | 0.125 | 0.048 | 0.119 | 0.054 |
| 4 | 0.647 | 0.570 | 0.132 | 0.189 | 83.455 | 33.821 | 65.603 | 28.389 | 0.117 | 0.045 | 0.079 | 0.046 |
| 5 | 0.652 | 0.628 | 0.127 | 0.161 | 86.391 | 41.412 | 52.239 | 41.645 | 0.162 | 0.052 | 0.100 | 0.048 |
| 6 | 0.641 | 0.589 | 0.089 | 0.193 | 86.314 | 35.767 | 73.863 | 43.355 | 0.178 | 0.045 | 0.126 | 0.051 |
| 7 | 0.636 | 0.615 | 0.119 | 0.190 | 85.209 | 25.563 | 75.789 | 26.788 | 0.165 | 0.039 | 0.122 | 0.036 |
| 8 | 0.627 | 0.553 | 0.146 | 0.188 | 59.652 | 29.889 | 38.877 | 23.420 | 0.131 | 0.044 | 0.110 | 0.041 |
| 9 | 0.653 | 0.644 | 0.116 | 0.199 | 80.385 | 36.571 | 89.120 | 29.299 | 0.145 | 0.040 | 0.100 | 0.036 |
| 10 | 0.618 | 0.643 | 0.096 | 0.153 | 66.364 | 31.462 | 53.194 | 24.653 | 0.128 | 0.049 | 0.095 | 0.036 |
| 11 | 0.665 | 0.586 | 0.123 | 0.137 | 95.193 | 31.045 | 109.020 | 22.933 | 0.165 | 0.053 | 0.126 | 0.049 |
| 12 | 0.629 | 0.608 | 0.119 | 0.156 | 93.033 | 42.100 | 86.004 | 32.527 | 0.177 | 0.060 | 0.129 | 0.057 |
| 13 | 0.627 | 0.611 | 0.124 | 0.126 | 96.462 | 36.267 | 77.527 | 31.066 | 0.194 | 0.056 | 0.110 | 0.061 |
| 14 | 0.635 | 0.616 | 0.098 | 0.169 | 77.600 | 32.281 | 50.341 | 37.162 | 0.168 | 0.047 | 0.102 | 0.042 |
| 15 | 0.657 | 0.591 | 0.124 | 0.175 | 86.400 | 42.692 | 52.435 | 47.113 | 0.157 | 0.047 | 0.120 | 0.052 |
| 16 | 0.643 | 0.634 | 0.096 | 0.165 | 113.083 | 37.172 | 107.830 | 31.304 | 0.203 | 0.056 | 0.135 | 0.051 |
| 17 | 0.647 | 0.610 | 0.127 | 0.156 | 81.879 | 32.976 | 67.533 | 34.301 | 0.164 | 0.051 | 0.106 | 0.051 |
| 18 | 0.622 | 0.594 | 0.115 | 0.149 | 65.529 | 25.769 | 43.408 | 20.117 | 0.129 | 0.045 | 0.068 | 0.045 |
| 19 | 0.630 | 0.605 | 0.104 | 0.196 | 97.721 | 38.075 | 89.398 | 30.737 | 0.192 | 0.059 | 0.119 | 0.048 |
| 20 | 0.625 | 0.547 | 0.138 | 0.149 | 65.355 | 44.000 | 61.241 | 45.981 | 0.137 | 0.068 | 0.118 | 0.069 |
| 21 | 0.618 | 0.529 | 0.131 | 0.237 | 88.500 | 49.667 | 77.735 | 34.070 | 0.152 | 0.053 | 0.103 | 0.028 |
| 22 | 0.653 | 0.556 | 0.158 | 0.129 | 56.808 | 69.200 | 40.750 | 59.198 | 0.119 | 0.089 | 0.092 | 0.065 |
| X | 0.653 | 0.588 | 0.143 | 0.179 | 74.163 | 32.346 | 83.889 | 43.351 | 0.143 | 0.040 | 0.129 | 0.042 |

**Figure 8:**
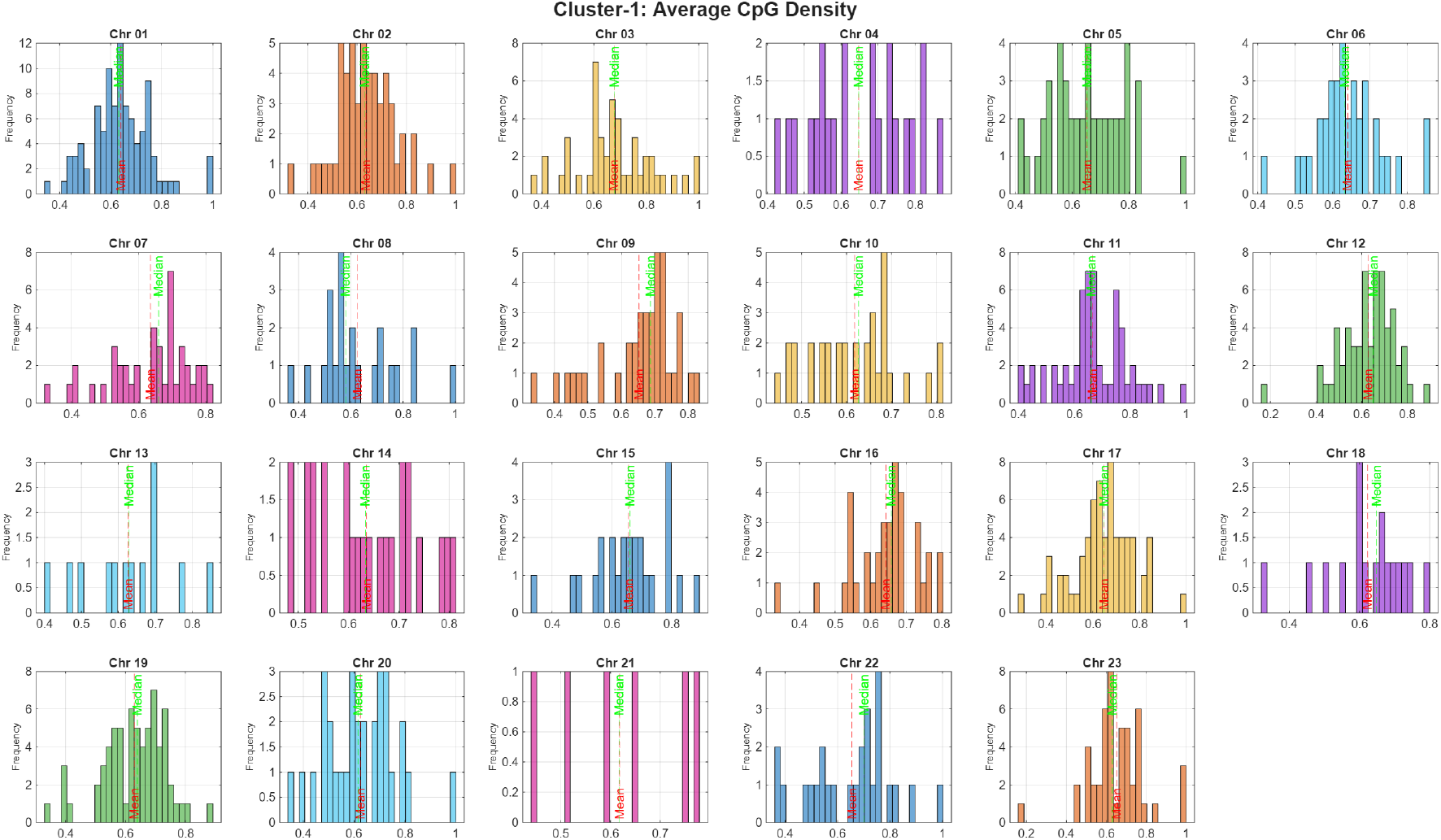
Histogram plots showing the distribution of average CpG density across human essential proteins for each human chromosome (1–23) from Cluster-1. Each subplot represents the frequency of CpG density values within a chromosome, highlighting intra-chromosomal variability and potential clustering patterns. Note that Chr 23 stands for chromosome X.

**Figure 9:**
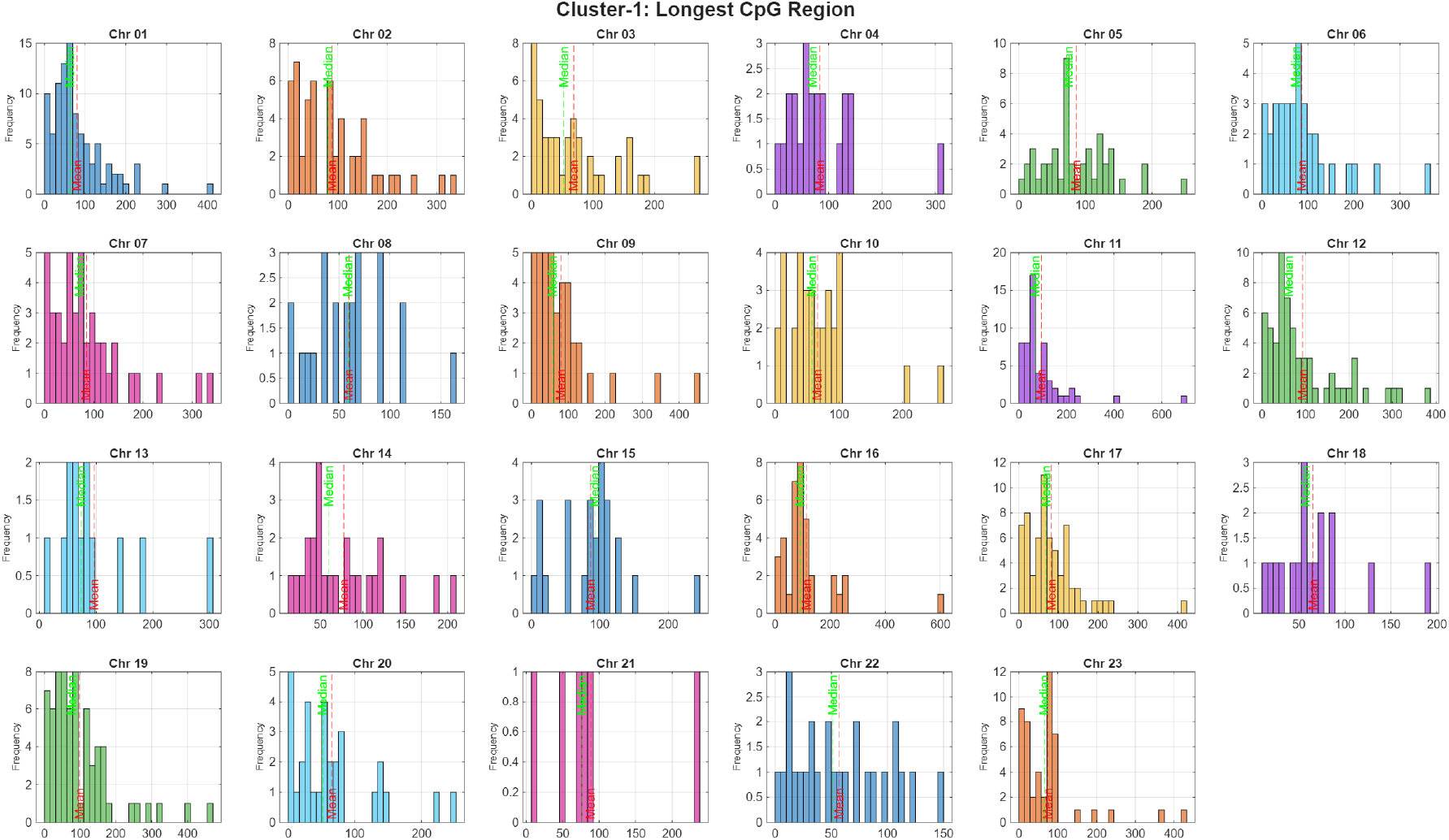
Histogram plots depicting the distribution of the longest CpG-induced amino acid region per human essential proteins across chromosomes 1–22 and X from Cluster-1. These plots reveal the frequency and spread of CpG-induced amino acid region lengths, with some chromosomes exhibiting broader or skewed distributions. Note that Chr 23 stands for chromosome X.

**Figure 10:**
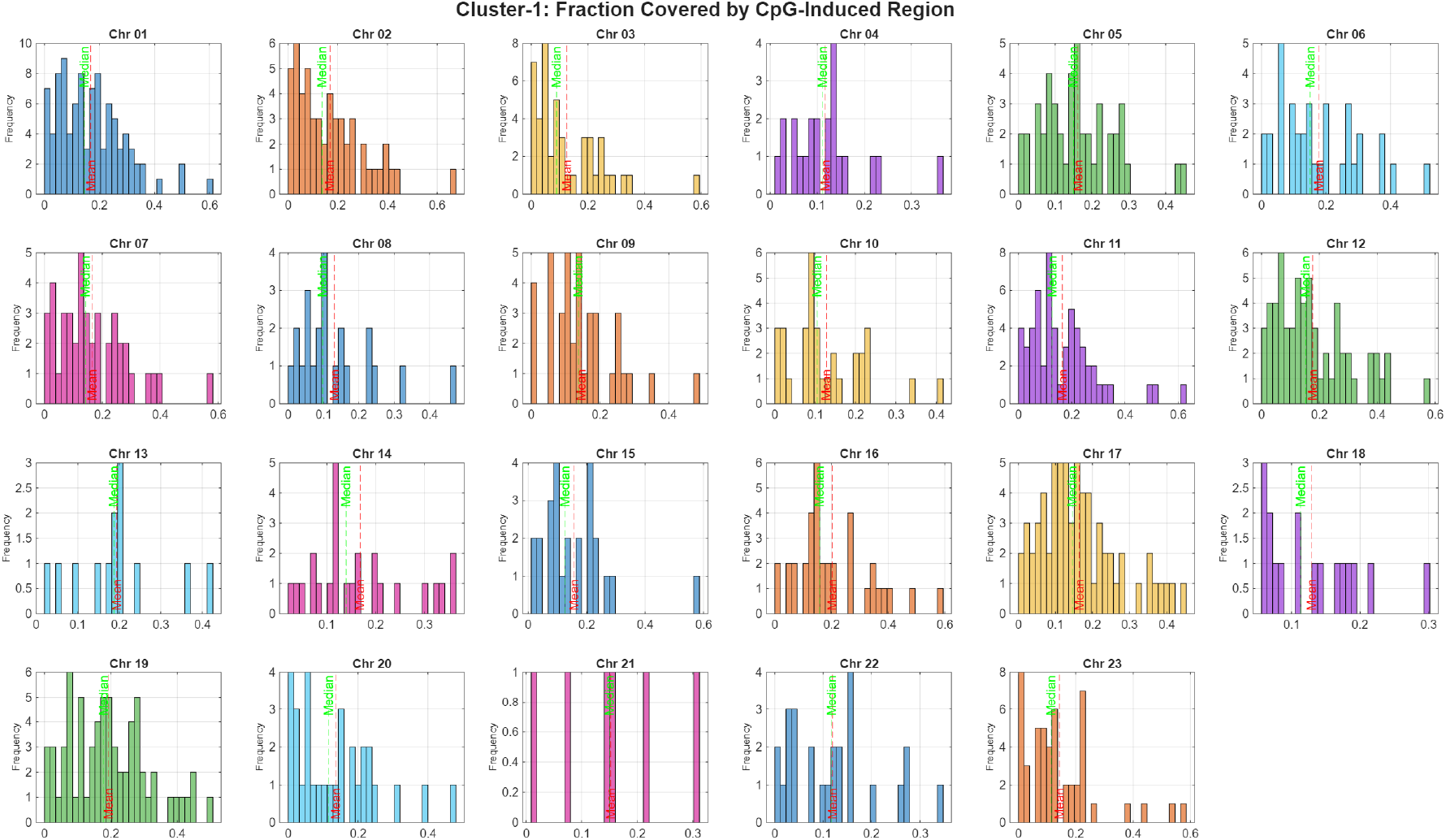
Histogram plots illustrating the distribution of CpG-induced amino acid region coverage across all human essential proteins per chromosome from Cluster-1. This metric indicates the proportion of each sequence occupied by CpG-induced amino acid region. Note that Chr 23 stands for chromosome X.

Across chromosomes, the same three CpG metrics were evaluated for Cluster-2 human essential proteins (Table 5). Histogram distributions of average CpG density ranged from 0.529 (Chromosome 21) to 0.644 (Chromosome 9), with elevated median values (≥0.669) in Chromosomes 3, 9, and 10 (Figure 11). Chromosomes 4, 8, 20, 21, and 22 showed lower median densities (≤0.554), indicating reduced CpG induced amino acid presence in the human essential proteins consisting CpG-induced regions (Table 5). The longest CpG-induced amino acid regions were most pronounced in Chromosome 22 (mean: 69.20), followed by Chromosomes 21 (49.67) and 20 (44.00), while Chromosomes 7, 8, 10, and 18 had shorter average regions (*<*32) (Figure 12, Table 5). Fractional coverage was highest in Chromosome 22 (mean and median: 0.089), followed by Chromosomes 20 (mean: 0.068) and 19 (mean: 0.059), with most other chromosomes exhibiting sparse CpG distribution (mean *<* 0.060) (Figure 13, Table 5). The highest variability was observed in Chromosome 13 (standard deviation: 0.061), while Chromosome 21 showed the most consistent coverage (standard deviation: 0.028) (Table 5). These histogram-based distributions suggest that CpG features in Cluster-2 proteins are generally more localized and less enriched compared to Cluster-1, though certain chromosomes retain elevated CpG characteristics.

**Figure 11:**
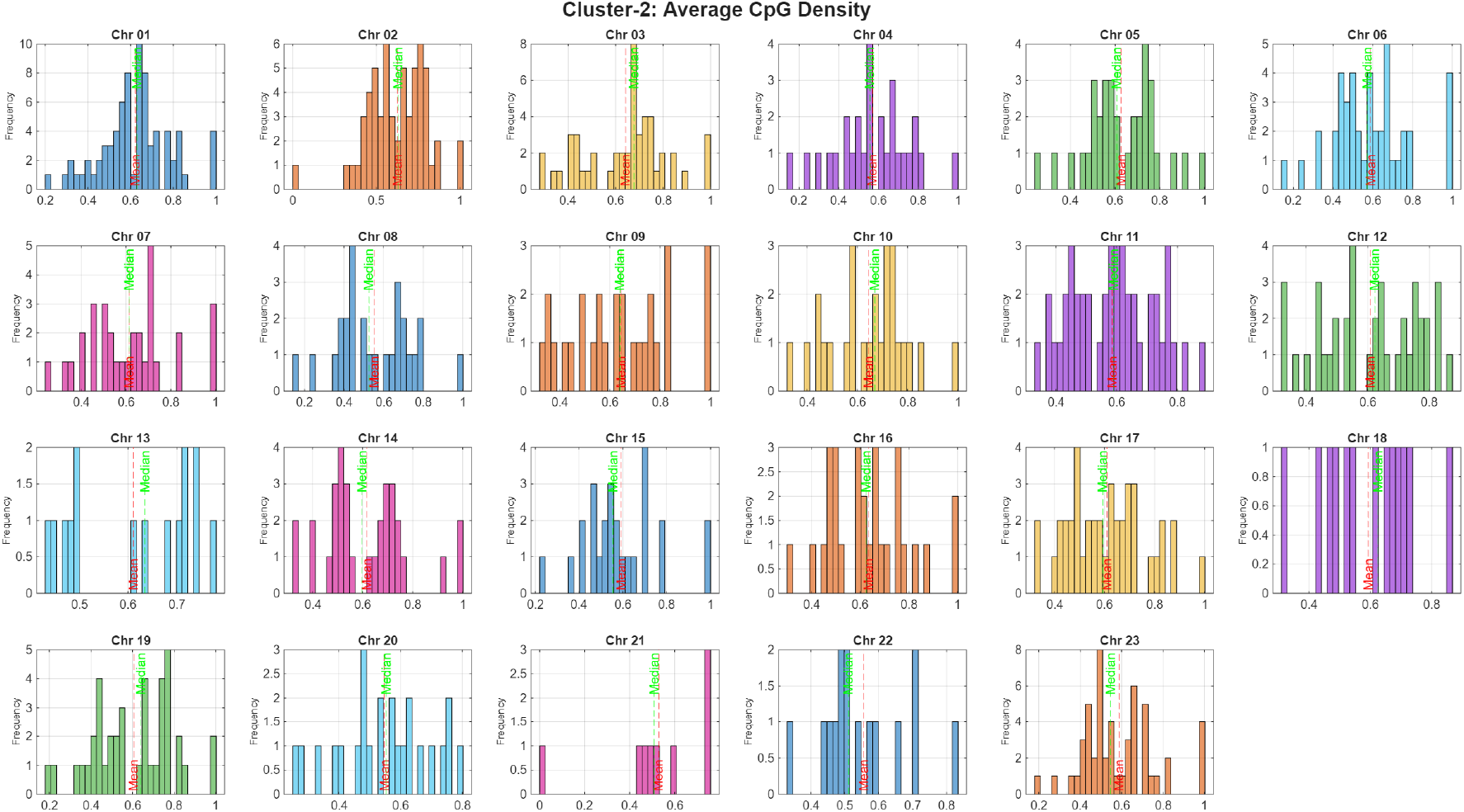
Histogram plots showing the distribution of average CpG density across human essential proteins for each human chromosome (1–22 and X) from Cluster-2. Each subplot represents the frequency of CpG density values within a chromosome, highlighting intra-chromosomal variability and potential clustering patterns. Note that Chr 23 stands for chromosome X.

**Figure 12:**
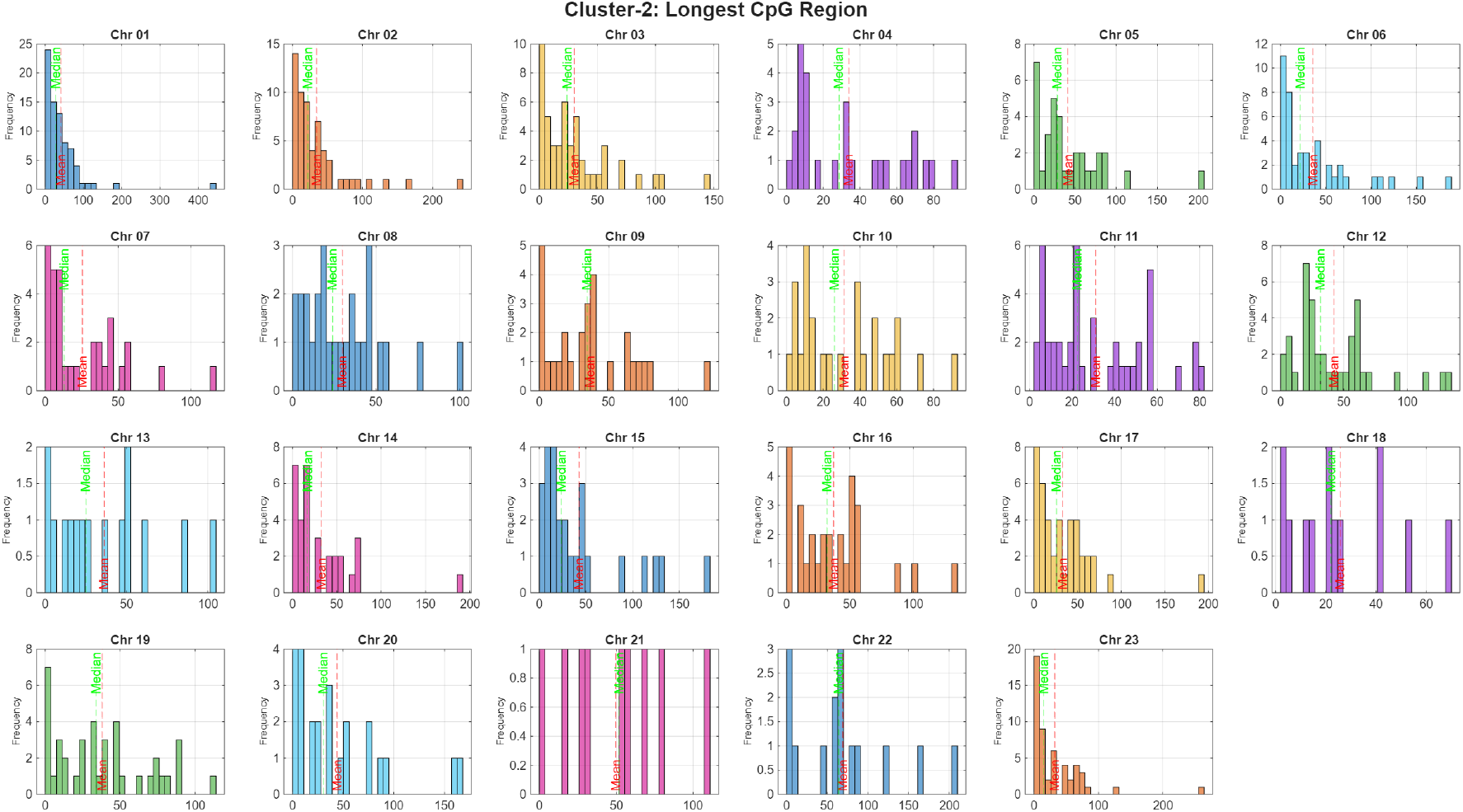
Histogram plots depicting the distribution of the longest CpG-induced amino acid region per human essential proteins across chromosomes 1–22 and X from Cluster-2. These plots reveal the frequency and spread of CpG-induced amino acid region lengths, with some chromosomes exhibiting broader or skewed distributions. Note that Chr 23 stands for chromosome X.

**Figure 13:**
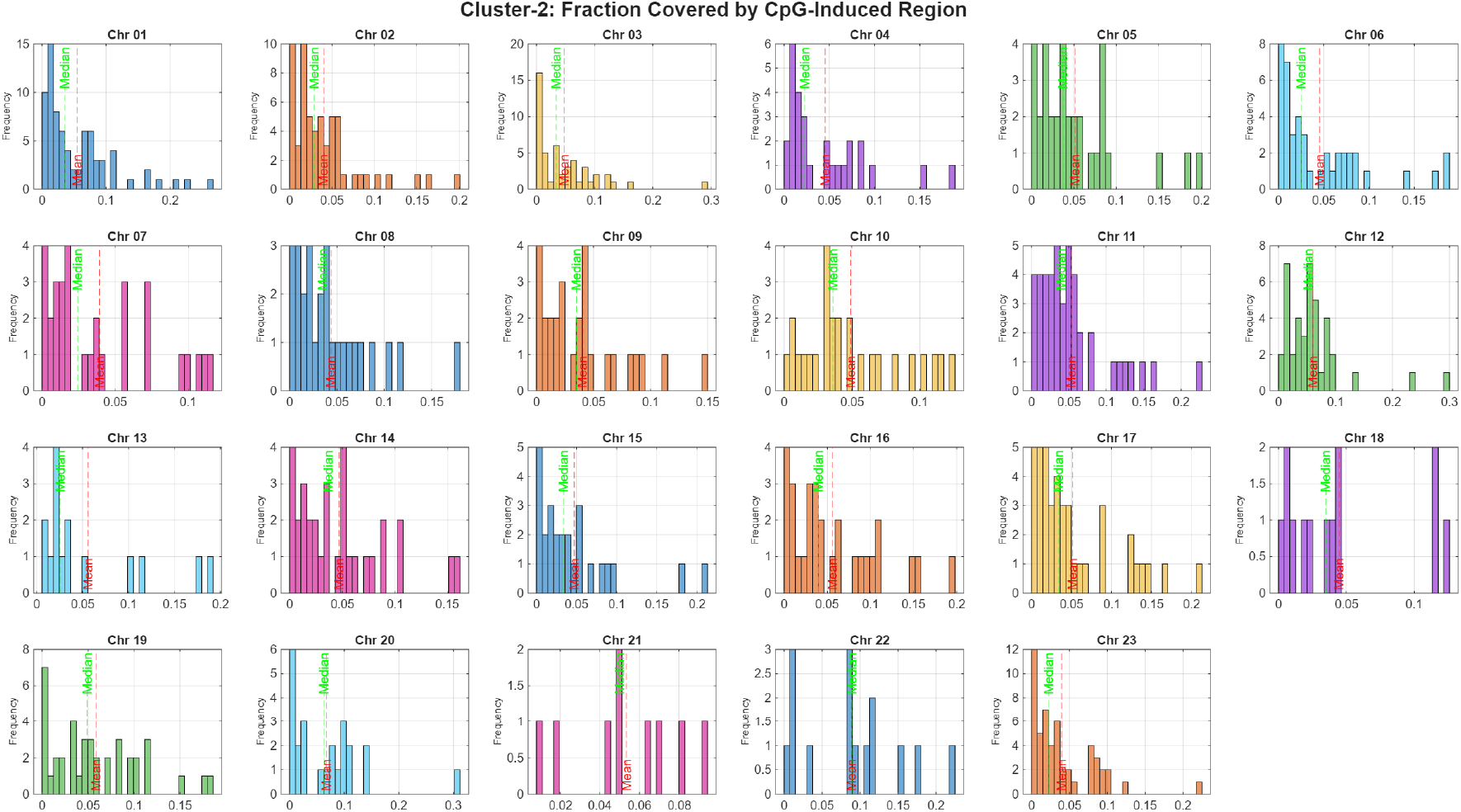
Histogram plots illustrating the distribution of CpG-induced amino acid region coverage across all human essential proteins per chromosome from Cluster-2. This metric indicates the proportion of each sequence occupied by CpG-induced amino acid region. Note that Chr 23 stands for chromosome X.

A comparative analysis of Cluster-1 and Cluster-2 proteins reveals distinct chromosomal architectures within CpG-induced amino acid regions. Cluster-1 proteins consistently exhibit higher average CpG density, longer CpG-induced amino acid regions, and greater fractional coverage across most chromosomes, reflecting a more CpG-enriched structural profile (Table 5). However, Cluster-2 proteins surpass Cluster-1 in specific contexts. Notably, Chromosome 10 shows higher CpG density in Cluster-2 (0.643 vs. 0.618), while Chromosomes 20, 21, and 22 exhibit longer CpG-induced amino acid regions, with Chromosome 22 reaching a mean of 69.20 compared to 56.81 in Cluster-1 (Figure 14). Additionally, Cluster-2 proteins on Chromosome 22 maintain the highest fractional coverage within the cluster (0.089), exceeding their Cluster-1 counterparts (Figure 14). These exceptions suggest that while CpG-induced region depletion is predominant in Cluster-2, certain chromosomes retain structural features that may support localized CpG-induced region induction or regulatory potential. Cluster-1 consistently shows higher mean values across most metrics, while Cluster-2 exhibits greater variability in segment properties (Figure 14).

**Figure 14:**
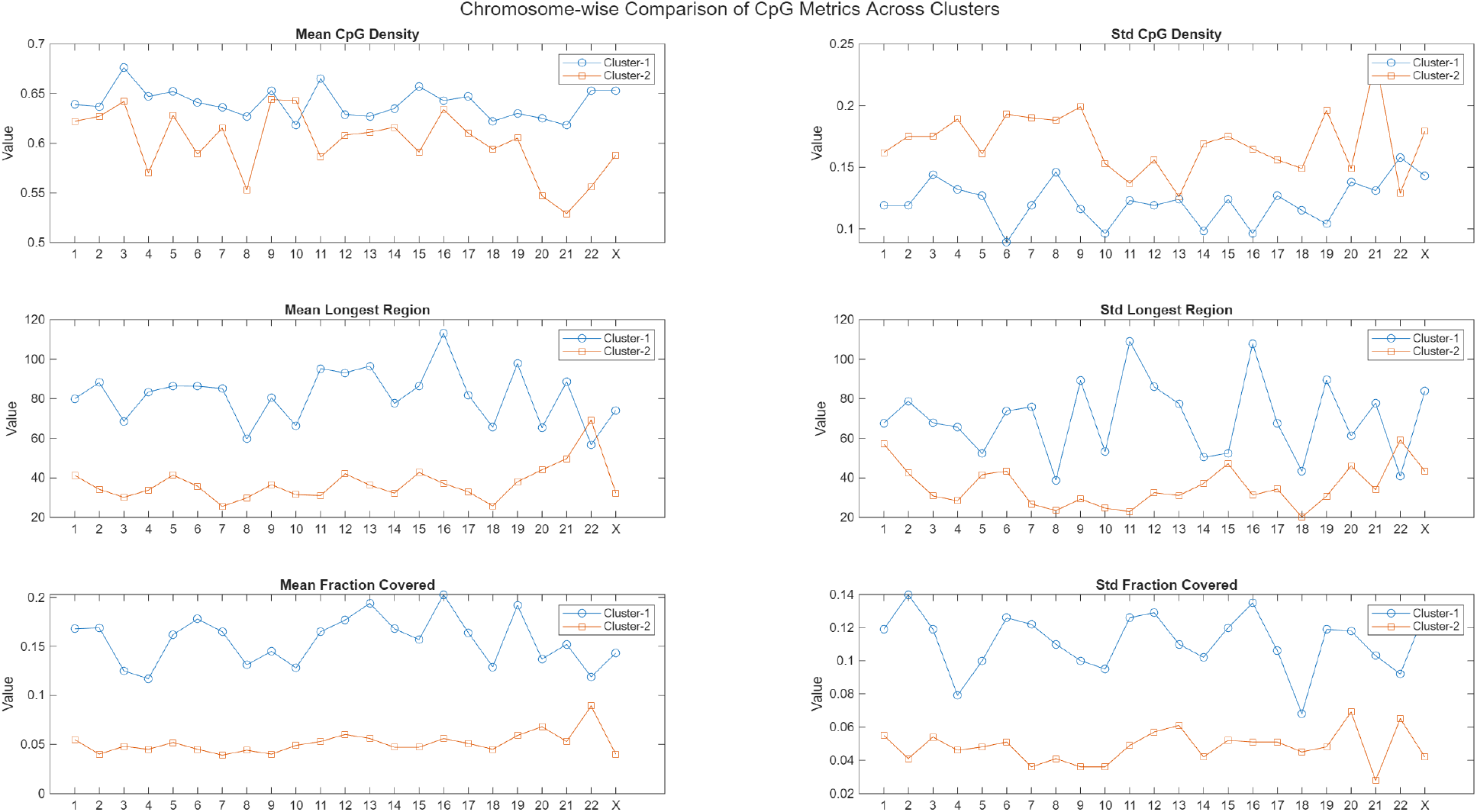
Chromosome-wise comparison of CpG-induced segment metrics between Cluster-1 and Cluster-2.

#### 4.4.2. Characteristics of the longest CpG-induced regions in human essential proteins

Proteins with pronounced polar enrichment were identified by extracting the longest CpG-induced segments from human essential genes exhibiting high PolarRatio values (≥ 0.8). A total of 16 proteins met this criterion, comprising 7 from Cluster-1 and 9 from Cluster-2, with segment lengths ranging from 10 to 114 amino acids. The highest PolarRatio was observed in *RBM25* (Cluster-2, Chromosome 14, PolarRatio = 1.0), followed by *GPATCH8* (Cluster-1, Chromosome 17, PolarRatio = 0.909) and *LUC7L3* (Cluster-2, Chromosome 17, PolarRatio = 0.882). These proteins represent top candidates for motif-specific annotation and comparative analysis, particularly in the context of CpG-induced sequence features and structural polarity (**Supplementary file–4**).

Conversely, to identify proteins with strong non-polar enrichment, we extracted CpG-induced segments with low PolarRatio values ( ≤ 0.2). This yielded 132 proteins, evenly distributed between Cluster-1 (*n* = 66) and Cluster-2 (*n* = 66). Segment lengths ranged from 10 to 413 amino acids, with the longest non-polar-rich segment found in *FMN2* (Cluster-1, Chromosome 1, 413 aa, PolarRatio = 0.1). The lowest PolarRatio values (0) were observed in *RBM22* (Cluster-1, Chromosome 5), *GDF11* (Cluster-2, Chromosome 12), and *PKM* (Cluster-2, Chromosome 15), indicating complete absence of polar residues in their CpG-induced amino acid regions. These essential proteins represent top candidates for hydrophobic motif analysis and contrast sharply with polar-rich profiles, suggesting distinct compositional biases and potential functional divergence (**Supplementary file–4**).

Histogram analysis of PolarRatio values across chromosomes revealed distinct cluster-specific trends. Cluster-1 proteins exhibited tighter distributions with elevated polar enrichment on several chromosomes, indicating consistent motif bias and reduced variability (Figure 15a). In contrast, Cluster-2 proteins showed broader and more heterogeneous distributions, with several chromosomes displaying flattened or bimodal patterns, suggesting greater diversity in polar residue composition. These contrasting profiles reflect potential structural or functional divergence linked to CpG-induced segment properties (Figure 15b).

**Figure 15:**
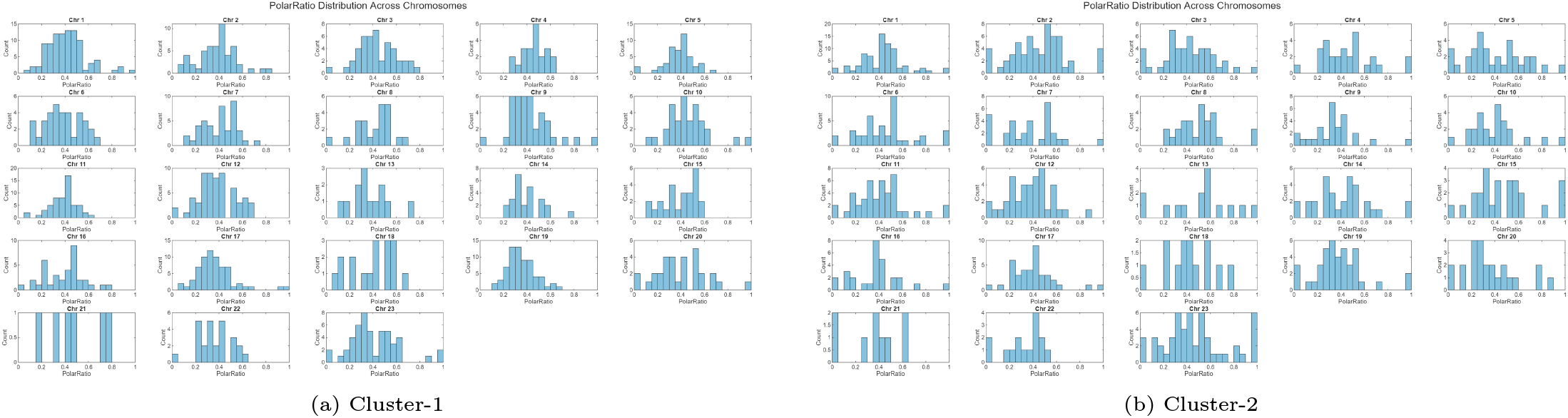
Distribution of polar residue ratios (PolarRatio) across chromosomes for Cluster-1 and Cluster-2 proteins. Each subplot shows chromosome-wise histograms of polar enrichment in longest CpG-induced regions. Note that Chr 23 stands for chromosome X.

To highlight proteins with elevated acidic residue content, we extracted the longest CpG-induced segments from human essential genes exhibiting high acidic enrichment (AcidicFrac ≥ 0.2). A total of 86 proteins satisfied this criterion, comprising 29 from Cluster-1 and 57 from Cluster-2, with segment lengths ranging from 10 to 115 amino acids. In Cluster-1, acidic enrichment was moderate, with values spanning 0.200–0.385. Notable examples included *PODXL2* (Chromosome 3, AcidicFrac = 0.385), *TBR1* (Chromosome 2, AcidicFrac = 0.286), and *ANKRD11* (Chromosome 16, length = 115 aa, AcidicFrac = 0.200) (**Supplementary file-4**).

In contrast, Cluster-2 displayed a broader range of acidic enrichment, with several proteins exceeding AcidicFrac = 0.4. The strongest enrichment was observed in *KIAA1468* (Chromosome 18, AcidicFrac = 0.462), *RBM25* (Chromosome 14, AcidicFrac = 0.444), and *PHF6* (Chromosome 23, AcidicFrac = 0.455). Additional highly enriched proteins included *WDFY3* (Chromosome 4, AcidicFrac = 0.389) and *DHX37* (Chromosome 12, AcidicFrac = 0.392). Chromosomes 1, 2, 7, 12, and 19 were particularly enriched in acidic-biased proteins within Cluster-2, suggesting chromosomal hotspots for acidic motif accumulation (**Supplementary file-4**).

Together, these findings indicate that while both clusters harbor acidic-rich CpG-induced regions, Cluster-2 proteins exhibit stronger and more variable acidic enrichment compared to Cluster-1. These subsets represent top candidates for motif-specific annotation and comparative analysis, particularly in the context of CpG-induced sequence features and charge-based structural preferences.

Histogram analysis of acidic fraction values across chromosomes revealed clear cluster-specific differences (Figure 16a, Figure 16b). Cluster-1 proteins generally exhibited narrower distributions centered around moderate acidic fractions (0.20–0.30), with only a few chromosomes (e.g., Chr 3 and Chr 7) displaying higher enrichment peaks. In contrast, Cluster-2 proteins showed broader and more heterogeneous distributions, with several chromosomes (e.g., Chr 7, Chr 12, Chr 14, Chr 18, and Chr 23) presenting pronounced peaks at higher acidic fractions ( ≥ 0.35). Notably, extreme enrichment was consistent with individual protein observations, including *KIAA1468* (Chr 18, AcidicFrac = 0.462), *RBM25* (Chr 14, AcidicFrac = 0.444), and *PHF6* (Chr 23, AcidicFrac = 0.455). These contrasting profiles indicate that while acidic enrichment is present in both clusters, Cluster-2 proteins display stronger and more variable acidic bias compared to Cluster-1. Such patterns suggest functional divergence and potential chromosomal hotspots for acidic motif accumulation (**Supplementary file-4**).

**Figure 16:**
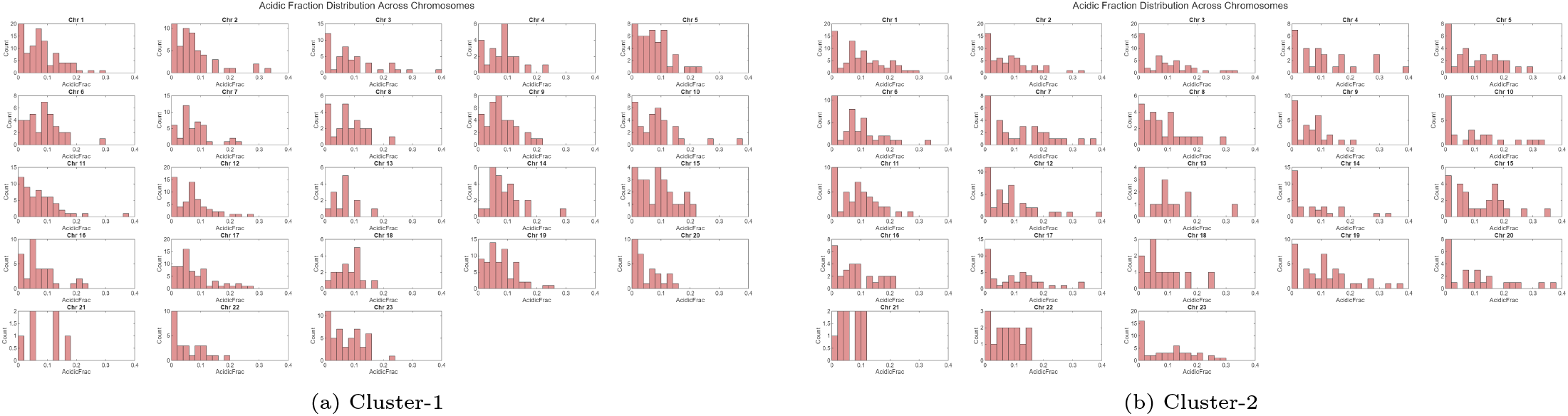
Distribution of acidic residue fractions (AcidicFrac) across chromosomes for Cluster-1 and Cluster-2 proteins. Histograms reflect D and E residue enrichment in CpG-induced regions. Note that Chr 23 stands for chromosome X.

To highlight proteins with elevated basic residue content, we extracted the longest CpG-induced segments from human essential genes exhibiting high BasicFrac values ( ≥ 0.2), reflecting enrichment in arginine (R), lysine (K), and histidine (H). A total of 289 proteins satisfied this criterion, comprising 146 from Cluster-1 and 143 from Cluster-2, with segment lengths ranging from 10 to 341 amino acids.

In Cluster-1, BasicFrac values ranged from 0.200 to 0.615. Several splicing regulators and RNA-binding proteins showed strong enrichment, including *SRSF11* (Chromosome 1, 152 aa, BasicFrac = 0.428), *PPIG* (Chromosome 2, 95 aa, BasicFrac = 0.432), and *U2SURP* (Chromosome 3, 74 aa, BasicFrac = 0.392). The highest enrichment was noted in *SON* (Chromosome 21, 236 aa, BasicFrac = 0.475) and *ZNRF3* (Chromosome 22, 30 aa, BasicFrac = 0.467), while *CPEB4* (Chromosome 5, 13 aa) exhibited an extreme BasicFrac of 0.615.

In Cluster-2, BasicFrac values extended up to 0.554, with several proteins exceeding 0.4. Representative examples included *PRPF38B* (Chromosome 1, 81 aa, BasicFrac = 0.420), *ETAA1* (Chromosome 2, 20 aa, BasicFrac = 0.450), *WHSC1L1* (Chromosome 8, 19 aa, BasicFrac = 0.421), and *DDX23* (Chromosome 12, 49 aa, BasicFrac = 0.408). The strongest enrichment was observed in *RSRC2* (Chromosome 12, 114 aa, BasicFrac = 0.535), *ARGLU1* (Chromosome 13, 52 aa, BasicFrac = 0.538), and *LUC7L3* (Chromosome 17, 90 aa, BasicFrac = 0.533), all surpassing the 0.5 threshold. These proteins represent top candidates for motif-specific annotation and comparative analysis, particularly in the context of CpG-induced sequence features and charge-driven structural or regulatory functions (**Supplementary file-4**).

Histogram analysis of BasicFrac values across chromosomes revealed distinct distributional patterns between clusters. Cluster-1 proteins showed moderately skewed distributions, with several chromosomes (e.g., Chr 1, Chr 3, Chr 7, and Chr 17) enriched in proteins with BasicFrac values between 0.30–0.45, consistent with the prevalence of splicing regulators and chromatin-associated proteins (Figure 17a). In contrast, Cluster-2 proteins exhibited broader and more heterogeneous distributions, with multiple chromosomes (e.g., Chr 2, Chr 8, Chr 12, Chr 13, and Chr 17) displaying pronounced peaks at higher BasicFrac values ( ≥ 0.4). Notably, extreme peaks in the histograms corresponded to proteins such as *RSRC2, ARGLU1*, and *LUC7L3*, which exceeded 0.5 (Figure 17b).

**Figure 17:**
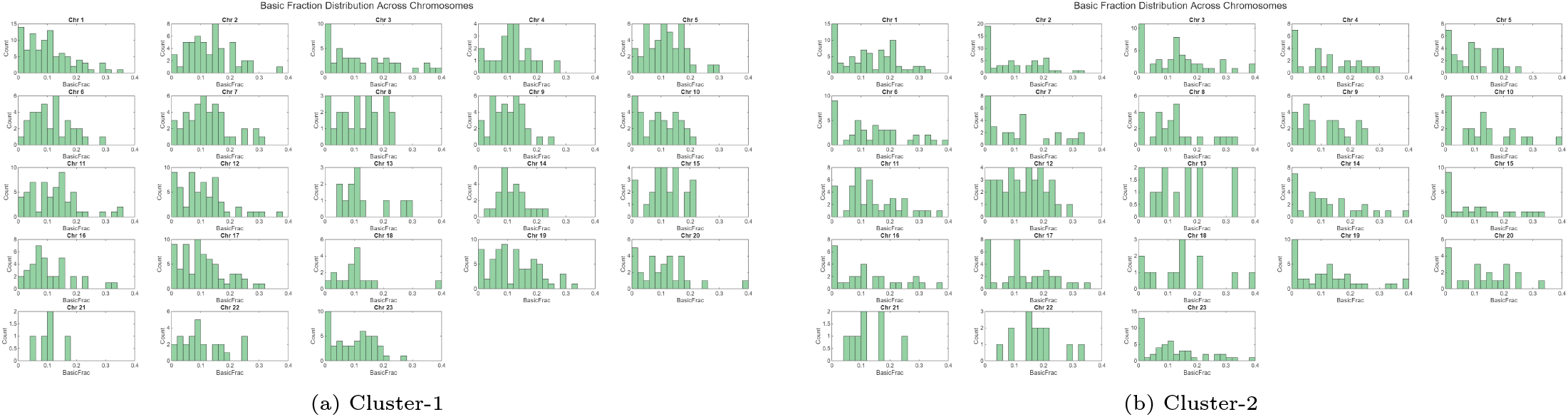
Distribution of basic residue fractions (BasicFrac) across chromosomes for Cluster-1 and Cluster-2 proteins. Histograms show R, K, and H residue enrichment patterns in CpG-induced regions. Note that Chr 23 stands for chromosome X.

These contrasting profiles indicate that while both clusters harbor basic-rich CpG-induced regions, Cluster-2 proteins display stronger and more variable basic enrichment compared to Cluster-1. Such patterns suggest functional divergence and potential chromosomal hotspots for basic motif accumulation.

#### 4.4.3. Residue composition varieties between clusters across chromosomes

Furthermore, to explore how residue composition varies between clusters, we compared the mean fractions of polar, acidic, and basic residues across chromosomes 1–22 and X for Cluster 1 and Cluster 2. These comparisons are visualized in Figure 18, which shows side-by-side trends for each metric. The chromosomes with the most pronounced differences are highlighted in Tables 6–8, with signed differences calculated as Δ = Cluster 2 − Cluster 1.

**Table 6:** Chromosomes with the largest signed differences in polar residue fraction.

| **Chromosome** | Cluster 1 | Cluster 2 | $\Delta_{\text{Polar}}$ |
| --- | --- | --- | --- |
| 13 | 0.379 | 0.501 | +0.122 |
| 21 | 0.473 | 0.348 | -0.125 |
| 8 | 0.405 | 0.503 | +0.098 |
| 9 | 0.428 | 0.339 | -0.089 |
| 2 | 0.381 | 0.451 | +0.070 |

**Table 7:** Chromosomes with the largest signed differences in acidic residue fraction.

| **Chromosome** | Cluster 1 | Cluster 2 | $\Delta_{\text{Acidic}}$ |
| --- | --- | --- | --- |
| 13 | 0.065 | 0.121 | +0.056 |
| 5 | 0.073 | 0.112 | +0.039 |
| 18 | 0.084 | 0.112 | +0.028 |
| 8 | 0.079 | 0.104 | +0.025 |
| 21 | 0.086 | 0.065 | -0.021 |

**Table 8:** Chromosomes with the largest signed differences in basic residue fraction.

| **Chromosome** | Cluster 1 | Cluster 2 | $\Delta_{\text{Basic}}$ |
| --- | --- | --- | --- |
| 10 | 0.091 | 0.190 | +0.099 |
| 22 | 0.118 | 0.206 | +0.088 |
| 6 | 0.113 | 0.178 | +0.065 |
| 7 | 0.122 | 0.181 | +0.059 |
| 17 | 0.106 | 0.181 | +0.075 |

**Figure 18:**
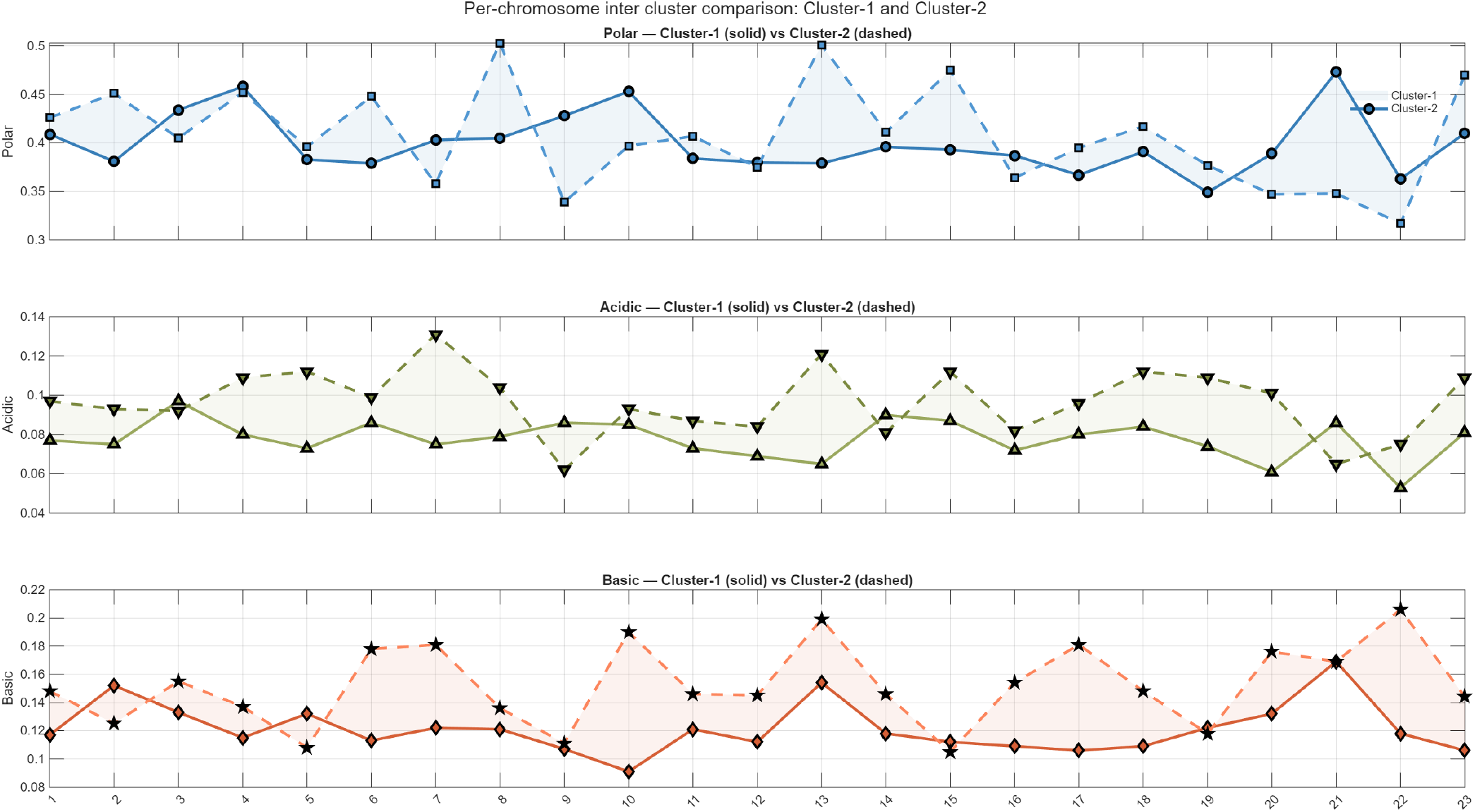
Cluster-wise comparison of residue composition across chromosomes. The figure displays three aligned line plots showing mean residue fractions for (A) polar, (B) acidic, and (C) basic residues across chromosomes 1–22 and X. Here 23 stands for the chromosome X.

#### Polar residues: opposing shifts across chromosomes

Polar composition varied substantially between clusters, but not in a uniform direction. Cluster 2 showed elevated polar ratios on chromosomes 13 (Δ = +0.122), 8 (+0.098), and 2 (+0.070). In contrast, Cluster 1 was enriched on chromosomes 21 (Δ = − 0.125), 9 ( − 0.089), and 7 ( − 0.045). These opposing shifts suggest that polar enrichment is localized to specific chromosomes rather than being a global feature of either cluster.

#### Acidic residues: modest and scattered changes

Compared to polar and basic residues, acidic fractions showed relatively small differences between clusters. The largest increase was observed on chromosome 13 (Δ = +0.056), followed by chromosomes 5 (+0.039) and 18 (+0.028). A few chromosomes, such as 21 and 14, showed slight decreases in Cluster 2. Overall, acidic content varied by less than ±0.06 across all chromosomes, suggesting limited discriminative power.

#### Basic residues: consistent enrichment in Cluster 2

Basic residue fraction showed the clearest and most consistent pattern across chromosomes. Cluster 2 was enriched on chromosome 10 (Δ = +0.099), chromosome 22 (+0.088), and chromosome 6 (+0.065). Additional gains were observed on chromosomes 7 and 17. These shifts suggest that basic composition is a strong and reliable marker of Cluster 2 identity.

### 4.5. Formation of subgroups of human essential proteins based on CpG-induced amino acid regions

Silhouette-based validation consistently supported a two-cluster structure across all chromosomes, enabling further subdivision into two subgroups per chromosome to capture intra-cluster heterogeneity (**Supplementary file-5 and Supplementary file-6**). Chromosome 01 exhibited the highest total protein count, with 96 proteins in Cluster 1 (Subgroup 1: 34, Subgroup 2: 62) and 76 proteins in Cluster 2 (Subgroup 1: 36, Subgroup 2: 40), reflecting substantial CpG diversity across all three metrics (Table 9). Chromosome 11 showed strong intra-cluster cohesion, with 54 proteins concentrated in Cluster 1 Subgroup 1 and a high silhouette score (0.848), consistent with its classification in the high CpG density tier. In contrast, Chromosome 12 displayed an inverse pattern, with 46 proteins in Cluster 1 Subgroup 2 and 37 in Cluster 2 Subgroup 1, suggesting bifurcation in CpG coverage and longest CpG-induced region metrics. Chromosome 17 was notably asymmetric, with 47 proteins in Cluster 1 Subgroup 1 and 28 in Cluster 2 Subgroup 2, consistent with moderate silhouette separation (0.527) and intermediate CpG density. Chromosomes 06 and 19 exhibited balanced distributions across clusters and subgroups, with silhouette scores of 0.614 and 0.568 respectively, and uniform CpG metric profiles. Chromosomes 13, 18, and 21 had lower overall protein counts and modest subgroup separation, yet retained silhouette scores above 0.68, indicating compact clustering despite reduced diversity; these chromosomes were classified in the low CpG density tier. Chromosome X demonstrated sex-linked asymmetry, with Cluster 1 dominated by Subgroup 1 (45 proteins) and Cluster 2 enriched in Subgroup 2 (17 proteins), consistent with its intermediate silhouette score (0.769) and mixed CpG metric stratification.

**Table 9:** Chromosome-wise distribution of proteins across Cluster 1 and Cluster 2 subgroups.

| Chromosome | Cluster-1 |  | Cluster-2 |  | Chromosome | Cluster-1 |  | Cluster-2 |  |
| --- | --- | --- | --- | --- | --- | --- | --- | --- | --- |
|  | *Subgroup-1* | *Subgroup-2* | *Subgroup-1* | *Subgroup-2* |  | *Subgroup-1* | *Subgroup-2* | *Subgroup-1* | *Subgroup-2* |
| Chromosome-01 | 34 | 62 | 36 | 40 | Chromosome-13 | 2 | 11 | 5 | 10 |
| Chromosome-02 | 38 | 15 | 55 | 4 | Chromosome-14 | 15 | 10 | 8 | 24 |
| Chromosome-03 | 33 | 13 | 40 | 9 | Chromosome-15 | 14 | 11 | 21 | 5 |
| Chromosome-04 | 18 | 4 | 12 | 16 | Chromosome-16 | 27 | 9 | 16 | 13 |
| Chromosome-05 | 24 | 22 | 24 | 10 | Chromosome-17 | 47 | 19 | 13 | 28 |
| Chromosome-06 | 7 | 28 | 16 | 27 | Chromosome-18 | 15 | 2 | 10 | 3 |
| Chromosome-07 | 30 | 13 | 20 | 12 | Chromosome-19 | 50 | 18 | 13 | 27 |
| Chromosome-08 | 16 | 7 | 8 | 19 | Chromosome-20 | 5 | 26 | 18 | 6 |
| Chromosome-09 | 30 | 9 | 19 | 9 | Chromosome-21 | 5 | 1 | 6 | 3 |
| Chromosome-10 | 24 | 9 | 15 | 11 | Chromosome-22 | 15 | 11 | 10 | 5 |
| Chromosome-11 | 54 | 3 | 30 | 14 | Chromosome-X | 45 | 4 | 35 | 17 |
| Chromosome-12 | 15 | 46 | 37 | 3 |  |  |  |  |  |

These findings underscore the utility of subgroup-level resolution in capturing CpG-driven structural and functional stratification across chromosomes, and highlight the interplay between clustering cohesion, CpG density tiers, and metric variability. Intra-cluster cohesion and inter-cluster separation are further visualized in three-dimensional scatter plots of CpG density, fraction covered, and longest CpG region (Figures 19), while subgroup-level distributions are summarized in Table 9.

**Figure 19:**
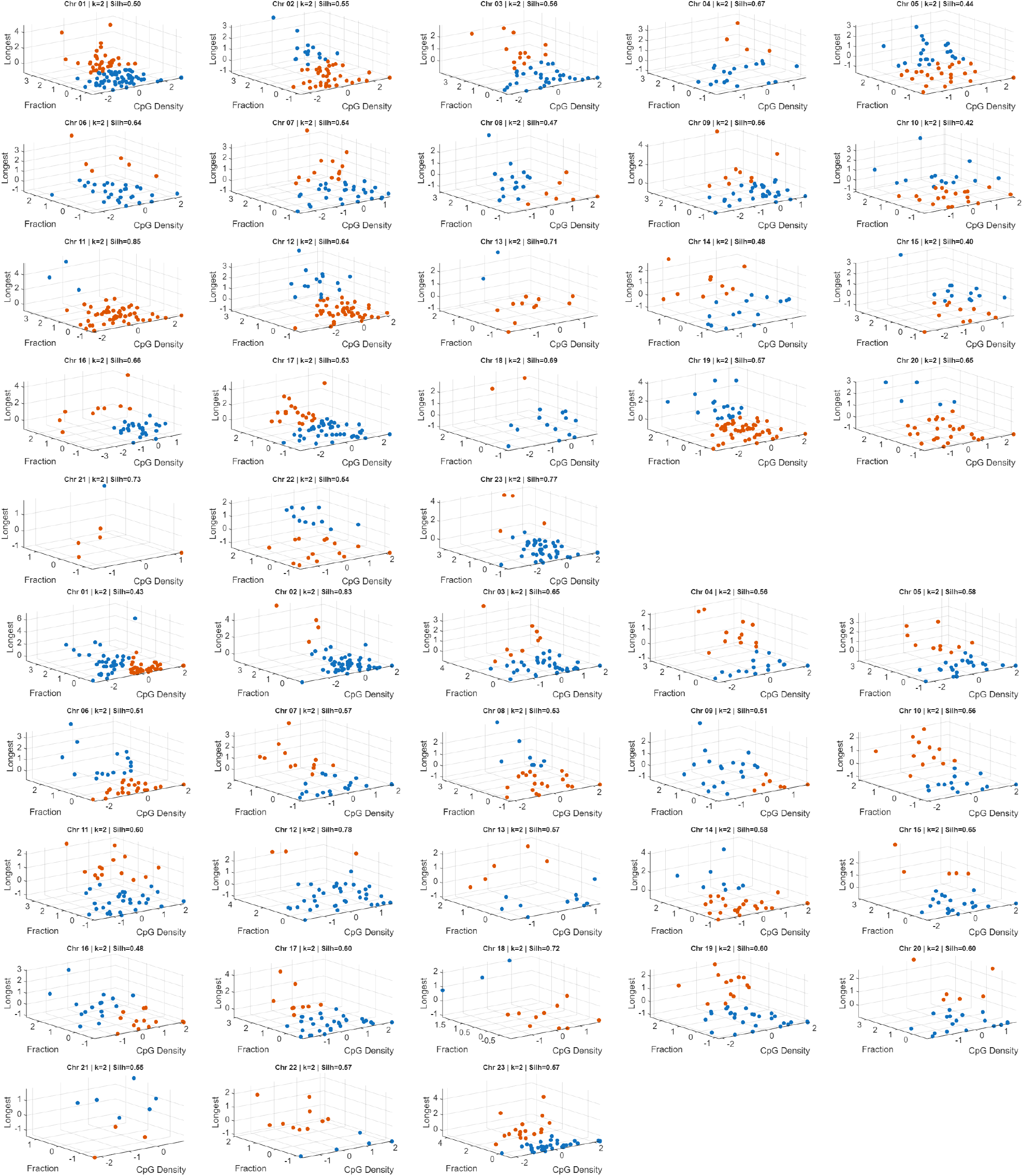
Three-dimensional scatter plots of protein clusters identified via k-means clustering using CpG density, fraction covered, and longest CpG region. Each point represents a protein, colored by cluster assignment. **Top:** Cluster 1. **Bottom:** Cluster 2. These visualizations highlight intra-cluster cohesion and inter-cluster separation across chromosomes. Here Chr 23 stands for chromosome X.

### 4.6. Chromosome-wise and subgroup-specific diversity in residue composition profiles of human essential proteins

Clustering of human essential proteins based exclusively on normalized percentages of polar, non-polar, acidic, and basic residues revealed distinct compositional landscapes across chromosomes and subgroups (Table 10). A detailed list of cluster assignments for each subgroup, chromosome, and cluster is provided in **Supplementary Files 7 and 8**. The optimal number of clusters for each chromosome and subgroup was determined using silhouette score maximization, ensuring biologically meaningful separation of residue-based profiles.

**Table 10:**
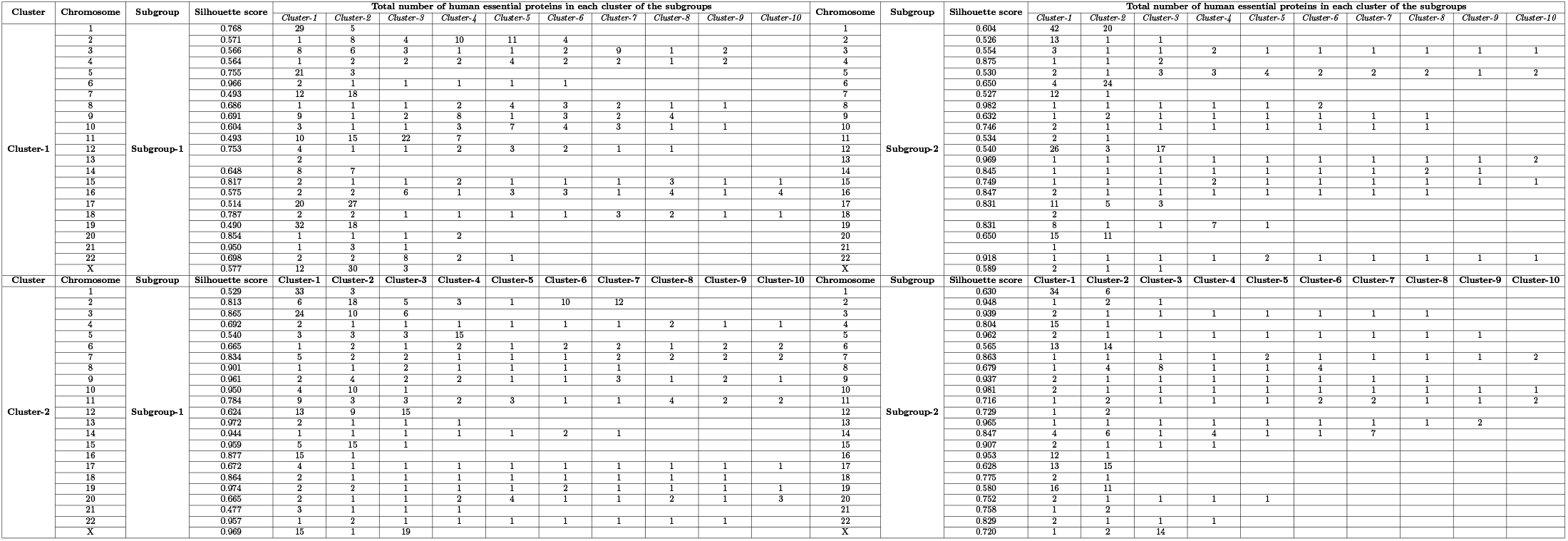
Chromosome-wise distribution of human essential proteins having CpG-induced amino acid regions across several optimal number of clusters within Subgroup-1 and Subgroup-2, shown separately for Cluster-1 and Cluster-2. Each cell represents the total number of human essential proteins assigned to a specific cluster on a given chromosome.

The resulting distributions highlight both conserved and divergent trends in amino acid composition. Several chromosomes—including 3, 4, 8, 9, 10, and 22—exhibited high cluster counts (8–10 clusters) with low per-cluster frequencies, often limited to one or two proteins (Table 10). This widespread dispersion suggests that essential proteins on these chromosomes possess highly variable combinations of physicochemical properties, with no dominant residue profile prevailing. Such diversity may reflect functional heterogeneity, structural adaptability, or differential evolutionary pressures acting on these regions.

In contrast, certain chromosomes demonstrated strong clustering biases. For example, Chromosome 19 in Subgroup-1 showed a pronounced accumulation of proteins in Cluster-1 (32 proteins), while Chromosome 17 in the same subgroup was heavily represented in Cluster-2 (27 proteins) (Table 10). These concentrated distributions imply conserved residue composition patterns—such as consistent polarity or charge profiles—potentially linked to shared structural motifs or functional domains among proteins encoded on these chromosomes.

Subgroup-specific shifts in cluster occupancy were also evident. Chromosome 6 transitioned from a dispersed multi-cluster profile in Subgroup-1 to a highly concentrated representation in Cluster-2 within Subgroup-2 (24 proteins), suggesting subgroup-dependent reorganization of residue composition (Table 10). Similar transitions were observed in Chromosomes 12 and X, where dominant clusters varied markedly between subgroups. These shifts may reflect differential expression, post-translational modification potential, or regulatory divergence between subgroups.

Chromosomes such as 5, 11, and 14 displayed moderate representation across multiple clusters, indicating balanced compositional diversity. This pattern may correspond to proteins with intermediate polarity and charge distributions, supporting multifunctionality or broad interaction potential. Conversely, chromosomes with fewer essential proteins, such as 21 and Y, consistently formed fewer clusters with minimal dispersion, aligning with expectations of limited compositional variability due to smaller sample sizes.

The X chromosome exhibited notable subgroup-specific polarization. In Subgroup-1, Cluster-2 contained 30 proteins, whereas in Subgroup-2, Cluster-3 was dominant with 14 proteins (Table 10). This divergence may reflect sex-linked specialization in residue composition, potentially influencing protein stability, localization, or interaction specificity.

Finally, saturation of cluster occupancy was observed in select chromosomes, such as Chromosome 3 in Subgroup-1, which populated all ten clusters (Table 10). This maximal spread underscores the extensive diversity in residue composition and may indicate high evolutionary plasticity or functional breadth among the encoded essential proteins.

#### Case Studies: Compositional clustering and functional stratification of chromosomally localized essential proteins

Among the clustering outcomes presented in Table 10, two representative case studies were selected to examine the structural and functional unity or diversity of proteins in relation to their respective cluster assignments.

##### Case Study 1

A subset of human essential proteins assigned to Subgroup_2 within Cluster 1 and located on chromosome 13 was selected for detailed examination of their functional roles.

A set of eleven essential human proteins—*PAN3, FOXO1, TSC22D1, KLF12, DACH1, RBM26, SLAIN1, SPRY2, MBNL2, ARHGEF7*, and *MYO16* —was subjected to compositional profiling based on the percentage of polar, acidic, and basic residues. k-means clustering of these profiles yielded ten distinct optimal number clusters with Silhouette score 0.969, with only one cluster (Cluster 10) containing two proteins (*FOXO1* and *SLAIN1*), while the remaining proteins were uniquely assigned to separate clusters (Table 10).

The clustering outcome reflects a compositional logic that aligns with the proteins’ structural domains and functional roles. Proteins enriched in polar and basic residues—such as *FOXO1* and *SLAIN1* —were co-clustered, consistent with their roles in transcriptional regulation and cytoskeletal dynamics, respectively [50, 51]. Both proteins exhibit nuclear or cytoplasmic localization and rely on electrostatic interactions for DNA binding or microtubule association [50, 51].

Proteins with elevated acidic residue content, such as *SPRY2* (Cluster 1), were separated due to their membrane-associated signaling roles, where acidic patches facilitate protein-protein interactions and post-translational modifications [52]. Similarly, *MYO16* (Cluster 5), a motor protein involved in neuronal development, was distinguished by its balanced acidic and polar composition, supporting flexible domain interfaces required for cytoskeletal remodeling [53, 54].

RNA-binding proteins such as *RBM26* (Cluster 3) and *MBNL2* (Cluster 2) were assigned to distinct clusters due to differences in their basic-to-acidic residue ratios. *RBM26*, involved in mRNA splicing, exhibits a basic-rich profile conducive to RNA backbone interaction, while *MBNL2*, containing zinc finger motifs, displays a more balanced composition suited for alternative splicing regulation and nuclear-cytoplasmic shuttling [55, 56, 57].

Transcription factors including *KLF12* (Cluster 9), *DACH1* (Cluster 4), and *TSC22D1* (Cluster 6) were separated based on their unique compositional signatures. *KLF12* and *DACH1* are basic-rich, reflecting their DNA-binding and nuclear localization functions, whereas *TSC22D1* contains a leucine zipper motif and exhibits a balanced acidic/basic profile, facilitating dimerization and transcriptional repression [58, 59, 60].

*ARHGEF7* (Cluster 8), a Rho guanine nucleotide exchange factor, was distinguished by its polar and acidic composition, consistent with its role in actin cytoskeleton regulation and small GTPase interaction [61]. *PAN3* (Cluster 7), a component of the mRNA decay complex, was clustered separately due to its polar and basic-rich profile, supporting RNA binding and deadenylation activity [62].

Overall, the clustering based on residue composition successfully stratified proteins according to their structural domains and functional specializations. The distinct compositional profiles reflect evolutionary pressures shaping residue usage to optimize protein behavior, localization, and interaction potential. This approach provides a scalable framework for functional inference in compositional proteomics.

##### Case Study 2

A subset of human essential proteins from Subgroup_2, assigned to Cluster 2 and located on chromosome X, was examined in detail to elucidate their structural and functional roles.

Seventeen essential human proteins located on chromosome X and classified under Subgroup_2 were subjected to compositional clustering based on the percentage of polar, acidic, and basic residues. The resulting stratification yielded three distinct clusters: Cluster 1 (ZRSR2), Cluster 2 (ADGRG2, MBTPS2), and Cluster 3 (15 remaining proteins; Table 10). Literature-supported analysis confirms that these clusters reflect meaningful structural and functional groupings.

Cluster 3 comprises proteins with moderate-to-high polar and basic residue content, consistent with roles in transcriptional regulation, RNA processing, cytoskeletal remodeling, and ubiquitin-mediated signaling. For example, *TBL1X* functions as a transcriptional co-regulator via WD40-repeat domains; *RBM10* regulates alternative splicing through RNA-binding motifs; *WAS* and *DIAPH2* modulate actin cytoskeleton dynamics; and *HUWE1* and *CUL4B* participate in ubiquitin ligase complexes [63, 64, 65, 66, 67, 68]. Proteins such as *PHF8* (histone demethylase), *OPHN1* (Rho-GTPase signaling), and *TENM1* (neuronal adhesion) further illustrate the functional breadth unified by shared compositional features [69, 70, 71]. The prevalence of polar and basic residues supports nuclear localization, dynamic protein-protein interactions, and modular domain architecture.

Cluster 2 includes *ADGRG2* and *MBTPS2*, both membrane-associated proteins with elevated acidic residue proportions. *ADGRG2*, a G protein-coupled receptor, mediates epithelial ion transport and reproductive signaling, while *MBTPS2* is a membrane-bound protease involved in cholesterol regulation and ER stress response [72, 73]. Their acidic-rich profiles facilitate extracellular ligand recognition, transmembrane stability, and proteolytic activity, indicating functional coherence within this cluster [72, 73].

Cluster 1 contains the RNA splicing factor *ZRSR2*, which is essential for U12-type intron recognition. Its distinct basic-rich composition supports spliceosome assembly and RNA backbone interaction, structurally separating it from the other clusters [74, 75].

Taken together, the clustering reveals both unity and diversity: Cluster 3 proteins exhibit compositional and structural cohesion despite functional heterogeneity, while Clusters 1 and 2 represent specialized roles with distinct residue profiles. These findings support the hypothesis that amino acid composition serves as a proxy for structural modularity and functional specialization among chromosomally co-localized proteins.

The comparative analysis of essential human proteins from chromosome 13 (Case Study 1) and chromosome X (Case Study 2) demonstrates that compositional clustering based on polar, acidic, and basic residue percentages effectively captures both structural modularity and functional specialization. In both cases, proteins with similar compositional profiles were grouped into coherent clusters that aligned with known biological roles, domain architectures, and cellular localizations.

Clustered proteins enriched in polar and basic residues consistently exhibited nuclear or cytoplasmic localization and were involved in transcriptional regulation, RNA processing, and cytoskeletal dynamics. These features were evident in proteins such as *FOXO1, SLAIN1, RBM10*, and *DIAPH2*, which rely on electrostatic interactions and modular domains for their activity. Conversely, proteins with elevated acidic content—such as *SPRY2, MYO16, ADGRG2*, and *MBTPS2* —were associated with membrane signaling, proteolytic regulation, and extracellular ligand recognition, reflecting distinct structural demands.

The presence of singleton clusters (e.g., *ZRSR2* in Cluster 1 and *FOXO1* /*SLAIN1* in Cluster 10) further underscores the specificity of compositional signatures in defining unique functional niches. These findings affirm that residue-level compositional metrics serve as robust predictors of protein behavior and domain organization, offering a scalable framework for functional inference across genomic contexts.

Together, the case studies validate the utility of compositional clustering in proteomic stratification, revealing both conserved and divergent structural themes among chromosomally co-localized proteins.

### 4.7. CpG island-associated CG distribution in human essential genes

Genes with lower exon overlap fractions were higher in number compared to those with higher overlap values (Figure 20A). Bins with an exon overlap fraction of 0.5 or above exhibited a markedly higher CGI-associated CG density than bins containing genes with lower overlap fractions. In contrast, the density of non-CGI-associated CGs remained consistently low across genes, regardless of their level of CpG island association (Figure 20A).

**Figure 20:**
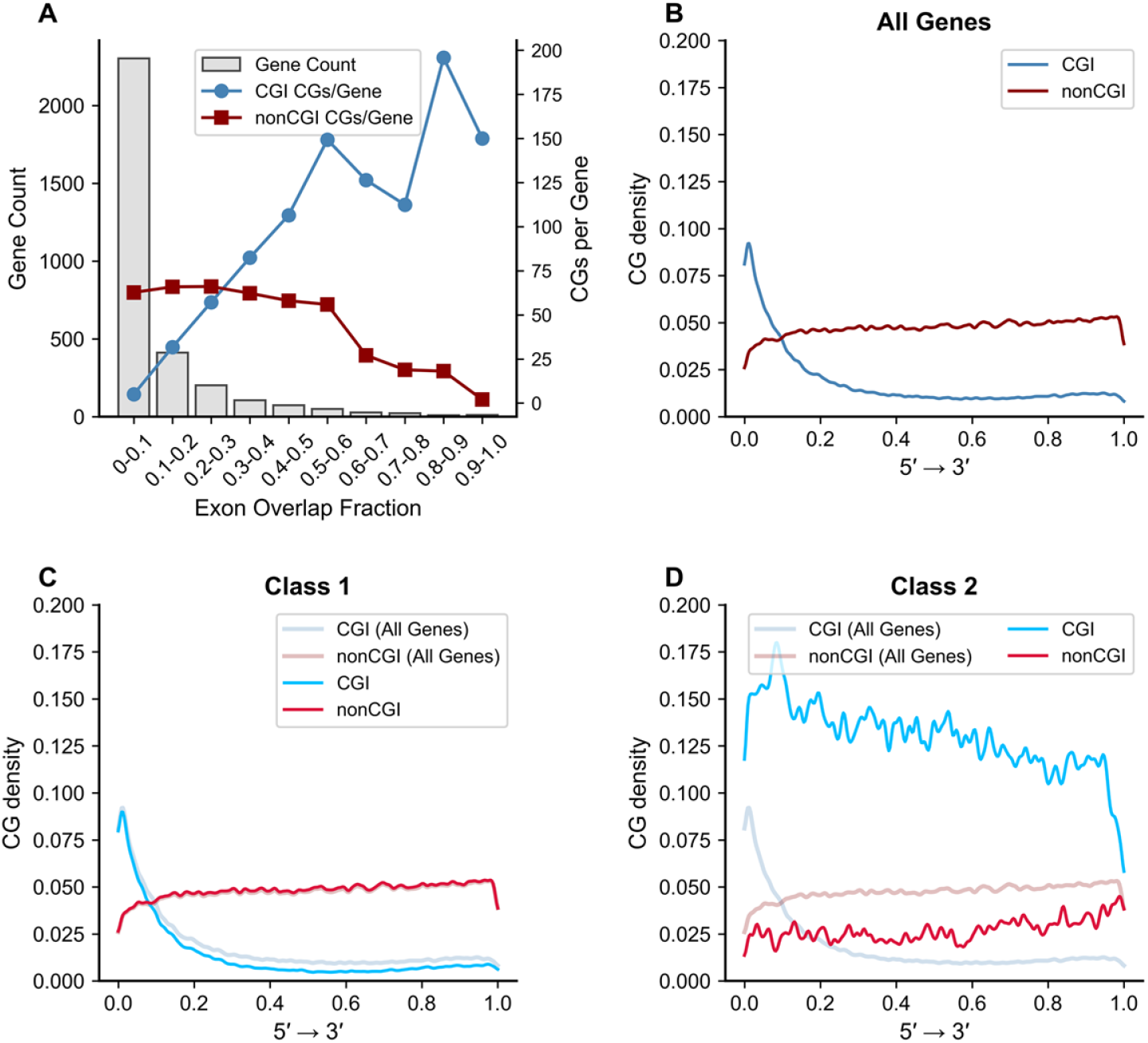
CpG distribution and density analyses across essential genes. A) Histogram of exon overlap fractions. Each bin represents the range of essential genes whose exonic regions overlap with CGI regions. The plot contains twin Y-axes: the left Y-axis indicates the number of genes in each bin, while the right Y-axis shows the CGI- and non-CGI-associated CG/gene count ratios calculated for all genes within each bin. B) CG density curves for CGI (blue) and non-CGI (red) regions plotted in the 5^*′*^ → 3^*′*^ direction for all genes across the exon overlap fraction bins. C) CG density curves for Class 1 (0-0.5 exon overlap fraction) shown in dark colors, overlaid with faint curves representing all essential genes (0-1.0 exon overlap fraction). D) CG density curves for Class 2 (0.5-1.0 exon overlap fraction) plotted together with faint background curves representing all genes across all overlap bins.

CG density curves plotted for all essential genes in the 5^*′*^ → 3^*′*^ direction clearly show that CGI-associated exon regions of most essential genes are enriched toward the 5^*′*^ end rather than the 3^*′*^ end of the genome (Figure 20B). In contrast, non-CGI-associated CG density remains uniformly low across nearly all regions. These observations indicate that CGI-associated CGs in essential genes are preferentially distributed toward the 5^*′*^ regions.

To further investigate the distribution of CGs from both CGI- and non-CGI-associated regions across essential genes, the genes were divided into two classes. Class-1 consisted of essential genes with an exon overlap fraction between 0 and 0.5, whereas Class-2 included genes with exon overlap fractions between 0.5 and 1. This classification was based on the CGI and non-CGI CG count fractions per gene obtained in Figure 20A, where a prominent increase in CGI-associated CG counts was observed beginning at the 0.5 exon-overlap bin.

The CG density curves for Class-1 displayed the same overall distribution pattern for both CGI and non-CGI regions as observed for all genes combined across all bins (Figure 20C). This indicates that essential genes with low exon overlap fractions still have the majority of their CGI-associated CGs concentrated toward the 5^*′*^ region of the genome, whereas the non-CGI-associated CGs, although fewer, are distributed throughout the gene body.

In Class-2 (Figure 20D), essential genes exhibited a very high and relatively uniform distribution of CGI-associated CGs across the genome, with a slight enrichment toward the 5^*′*^ end. The non-CGI-associated CGs in this subgroup showed a very low and uniform distribution across the genomic regions.

A total of 1,524 essential protein-coding genes previously classified as CpG-depleted were further analysed to determine their exon-overlap categories based on CpG island (CGI) overlap. Of these, only 27 genes were assigned to Class 2 (high exon overlap with CGIs), whereas the remaining 1,497 genes belonged to Class 1 (low exon overlap with CGIs). This distribution indicates that most CpG-depleted proteins are encoded by Class-1 essential genes, while CpG-induced proteins are distributed across both classes. Thus, the CGI-overlap–based classification also supports the definitions of different protein types depending upon CG- and non-CG encoding codons: CpG-depleted proteins are largely devoid of CGIs, whereas CpG-induced proteins can exhibit either low or high fractions of CGI overlap within their coding exons.

Therefore, CGI-associated CpGs of CGI regions show enrichment toward the 5^*′*^ end across both classes, as well as in most essential genes of the human genome.

## 5. Discussion

This study presents a comprehensive framework for understanding the compositional architecture of human essential proteins through the lens of CpG-associated amino acid usage. By integrating clustering, chromosomal mapping, entropy profiling, and residue-level analysis, we demonstrate that amino acid composition—particularly among CpG-enriched residues—is a non-random, context-dependent feature shaped by genomic architecture, mutational bias, and functional constraints.

Hierarchical and k-means clustering based on five CpG-sensitive residues (Alanine, Arginine, Serine, Proline, Threonine) revealed two distinct compositional regimes among 3,222 proteins. Principal Component Analysis (PCA) supported this separation, with PC1 and PC2 capturing 62% of the total variance. The inverse relationship between Serine and Alanine, and opposing contributions from Proline and Threonine, reflect trade-offs in residue usage likely driven by epigenetic regulation and selective pressures.

To further quantify compositional diversity, Shannon entropy was computed for each protein sequence. Entropy distributions revealed a statistically significant shift in complexity between clusters: Cluster 1 proteins exhibited lower entropy (mean *µ*_1_ = 3.91, standard deviation *σ*_1_ = 0.11, *n*_1_ = 961), while Cluster 2 proteins showed higher entropy (mean *µ*_2_ = 4.05, *σ*_2_ = 0.07, *n*_2_ = 2253). A Wilcoxon rank-sum test confirmed the difference (*p* = 5. 50 × 10^−145^), indicating that CpG-driven clustering reflects not only residue composition but also underlying sequence variability. Kernel density estimates revealed broader, left-skewed distributions in Cluster 1 and sharper peaks in Cluster 2, suggesting functional divergence in sequence modularity and constraint.

Cluster 1 proteins exhibited elevated Serine and Proline content with greater compositional heterogeneity, suggesting functional specialization and responsiveness to CpG island dynamics. Cluster 2 proteins showed tighter distributions and reduced variability, indicative of structural stability and reduced susceptibility to CpG mutability. Correlation analysis revealed moderate inter-residue dependencies in Cluster 1 and minimal coupling in Cluster 2, reinforcing compositional constraint.

Chromosomal stratification revealed non-uniform distribution of CpG-driven regimes. Chromosomes 19 and 17 were enriched in Cluster 1 proteins, aligning with high gene density and regulatory complexity. Chromosome X showed a pronounced bias toward Cluster 2, possibly reflecting sex-linked epigenetic regulation and dosage compensation. Across all chromosomes, Cluster 1 was predominantly CpG-induced, while Cluster 2 was CpG-depleted, suggesting a bifurcation between dynamic regulatory proteins and stable housekeeping components.

Compositional analysis of CpG-induced regions revealed distinct biases in polar, non-polar, acidic, and basic residue enrichment. Polar-rich segments (PolarRatio ≥ 0.8) were rare but tightly distributed in Cluster 1, suggesting conserved motif bias. Cluster 2 proteins exhibited broader polar profiles, reflecting structural diversity. Non-polar-rich segments (PolarRatio ≤ 0.2) were abundant and evenly distributed, indicating roles in membrane association and folding. Acidic enrichment (AcidicFrac ≥ 0.2) was stronger and more variable in Cluster 2, with hotspots on chromosomes 7, 12, 14, 18, and X. Basic residues (BasicFrac ≥ 0.2) were most prevalent, with Cluster 2 consistently enriched across chromosomes, supporting nucleic acid binding and chromatin interaction.

Comparative residue analysis across chromosomes revealed nuanced patterns: polar and acidic residues showed localized shifts, while basic residues emerged as robust markers of Cluster 2 identity. Subgroup-level stratification added further resolution, with Silhouette-based validation supporting a two-subgroup structure across all chromosomes. Chromosome 11 showed strong cohesion in Cluster 1 Subgroup 1, while chromosome 12 exhibited divergent distributions, indicating regulatory bifurcation. Chromosome X revealed sex-linked asymmetry, with distinct subgroup dominance in each cluster. Even chromosomes with lower protein counts maintained compact clustering, reinforcing biological relevance.

Two case studies further validated the compositional framework. In Case Study 1, eleven proteins from chromosome 13 were optimally partitioned into ten clusters (Silhouette score 0.969), reflecting strong intra-cluster similarity and functional coherence. Proteins enriched in polar and basic residues (e.g., *FOXO1, SLAIN1*) were associated with transcriptional regulation and cytoskeletal dynamics, while acidic-rich proteins (e.g., *SPRY2, MYO16*) were linked to membrane signaling and motor activity [76]. RNA-binding proteins (*RBM26, MBNL2*) and transcription factors (*KLF12, DACH1, TSC22D1*) were separated based on distinct compositional signatures, supporting the hypothesis that residue usage encodes functional modularity [77, 60].

Case Study 2 examined seventeen proteins from chromosome X, revealing three compositional clusters. Cluster 3 (15 proteins) exhibited structural cohesion despite functional heterogeneity, unified by moderate-to-high polar and basic residue content. Cluster 2 included membrane-associated proteins (*ADGRG2, MBTPS2*) with elevated acidic profiles, while Cluster 1 contained a single RNA splicing factor (*ZRSR2*) with a distinct basic-rich signature. These findings reinforce the utility of compositional metrics in stratifying proteins by domain architecture and cellular localization.

Together, these results demonstrate that CpG-associated compositional regimes reflect evolutionary pressures, structural modularity, and functional specialization. Shannon entropy emerges as a complementary metric to residue-level clustering, capturing latent complexity and variability across protein subsets. Residue-level stratification remains a scalable and interpretable method for proteomic classification, particularly valuable when structural data are limited. The integration of compositional logic with chromosomal context offers new avenues for motif annotation, epigenetic modeling, and disease association studies.

Our CGI–exon–overlap–based classification of essential genes further substantiates the differences previously observed between CpG-depleted and CpG-induced protein sets. The majority of CpG-depleted proteins were encoded by Class-1 genes, which show minimal CGI overlap, whereas CpG-induced proteins were distributed across both classes, reflecting a variable range of CGI enrichment within their coding exons. These results support the robustness of the CGI-based classification and confirm that CpG-depleted proteins are genuinely lacking in CpG island features, while CpG-induced proteins exhibit lower to higher degrees of CGI overlap.

Importantly, despite these differences, both classes showed a conserved pattern in the positional distribution of CpGs, with CpG enrichment consistently observed toward the 5^*′*^ ends of most essential genes. This shared feature suggests a common regulatory architecture that may be maintained across essential gene categories irrespective of their CGI overlap class.

Future investigations should integrate methylation landscapes, transcriptomic profiles, and structural modeling to clarify the mechanistic underpinnings and biomedical significance of these compositional signatures. Extending the analytical framework to encompass disorder propensity, hydrophobicity gradients, and charge distribution could further sharpen resolution and strengthen predictive capacity in functional proteomics.

## Acknowledgments

The authors sincerely thank Debaleena Nawn for her valuable contributions to technical discussions and for providing insightful suggestions that helped shape the direction of this study. Her handcrafted design of the graphical abstract and schematic flow chart is especially appreciated, offering a visually compelling and conceptually precise representation of the study’s core framework.

## Author contributions statement

SSH, SS, and KH formulated the problem and designed the theoretical experiments. SSH, SS, and KH carried out the experiments and performed the analyses. SSH, SS, KH, and VNU drafted the initial manuscript. All authors contributed to reviewing and editing the manuscript. SSH and KH supervised the overall project. All authors reviewed, checked, and approved the final manuscript.

